# Neural differences in conflict monitoring and stimulus expectancy processes in experienced meditators are likely driven by enhanced attention

**DOI:** 10.1101/2024.12.23.630201

**Authors:** Aron T. Hill, Sung Wook Chung, Melanie Emonson, Andrew W. Corcoran, Bernadette M. Fitzgibbon, Paul B. Fitzgerald, Neil W. Bailey

## Abstract

**Objectives:** Mindfulness meditation has been linked to enhanced attention and executive function, likely resulting from practice-related effects on neural activity patterns. In this study, we used an event-related potential (ERP) paradigm to examine brain responses related to conflict monitoring and attention in experienced mindfulness meditators to better understand key factors driving meditation-related effects.

**Methods:** We measured electroencephalography-derived N2 and P3 ERPs reflecting conflict monitoring and attention processes from 35 meditators and 29 non-meditators across both an easy and a hard Go/Nogo task (50% Nogo and 25% Nogo stimuli, respectively).

**Results:** Meditators displayed distinct neural activity patterns compared to non-meditators, with enhanced N2 responses in fronto-midline electrodes following hard Nogo trials (*p*_FDR_ = 0.011, *np^2^* = 0.111). The fronto-midline N2 ERP was also larger following Nogo trials than Go trials, in the harder task condition, and was related to correct responses. Meditators also exhibited a more frontally distributed P3 ERP in the easy task compared to the hard task, while non-meditators showed a more frontally distributed P3 ERP in the hard task (*p*_FDR_ = 0.015, *np^2^* = 0.079).

**Conclusions:** Mindfulness meditation was associated with distinct topographical patterns of neural activity in the attention task, without corresponding increases in global neural activity amplitudes. These meditation-related effects appear to be driven by attention-specific mechanisms, despite the examined neural activity being associated with conflict monitoring and stimulus expectancy. Our findings suggest that the cognitive benefits of meditation may only emerge in tasks that actively engage targeted cognitive processes, such as sustained attention.

## Introduction

Mindfulness meditation is a mental training technique that typically involves the deliberate selection of an attentional focus, often to the breath or bodily sensations, and the constant direction and re-direction of attention to this focus with an attitude of nonjudgemental awareness (Crane et al., 2017; Van Dam et al., 2018). This practice has been associated with enhanced psychological well-being (Goyal et al., 2014; McClintock et al., 2019; Querstret et al., 2020) as well as improved cognitive performance across specific domains (Im et al., 2021; Sumantry & Stewart, 2021; Yakobi et al., 2021).

Two cognitive processes are commonly reported to be improved by mindfulness meditation: attention and executive control (a function which prioritises the processing of stimuli, cognitive processes and responses related to internal goals). Significant mindfulness-related improvements in accuracy (but not reaction time) are commonly reported from tasks that measure these constructs (Im et al., 2021; Sumantry & Stewart, 2021; Verhaeghen, 2021; Yakobi et al., 2021). Theoretical perspectives, supported by some empirical evidence, suggest that enhanced mindful attention may serve as a core mechanism underlying meditation’s positive effects on well-being. Specifically, by improving one’s ability to focus on the present moment, such attentional refinement may decrease time spent ruminating, worrying, or experiencing anxious thoughts (Baer, 2009; Coffey et al., 2010; Gu et al., 2015; Johannsen et al., 2022).

Mindfulness meditation has also been shown to modify the neural activity underlying attention and executive control. For example, functional magnetic resonance imaging (fMRI) indicates that meditation is associated with increased activation in brain regions related to attention and executive function, including the anterior cingulate cortex (ACC), posterior cingulate cortex (PCC), dorsolateral prefrontal cortex (DLPFC), and insula (Allen et al., 2012; Falcone & Jerram, 2018; Ganesan et al., 2022; Tang et al., 2010; Tomasino et al., 2016). Meta-analyses also indicate that focused attention meditation engages the brain’s executive control network, and that meditation experience impacts activity in attention-related brain regions (Falcone & Jerram, 2018; Ganesan et al., 2022). Additionally, research using electroencephalography (EEG) to explore meditation-related electrical brain activity patterns has shown differences between meditators and non-meditators in attention- and executive control-related neural activity (Andreu et al., 2019; Bailey et al., 2023a; Bailey et al., 2019a; Bailey et al., 2023e; Moore et al., 2012; Yoshida et al., 2020).

Although researchers have broadly concluded that mindfulness meditation improves attention and executive function, results remain inconsistent, with some studies reporting null results (Bailey et al., 2019b; Fucci et al., 2022; Payne et al., 2020). These null findings may partially reflect the influence of individual factors on meditation-related effects, for example, the extent of meditation experience, practice type, age, and baseline cognitive capacity (as higher baseline capacity might allow less room for improvement from mindfulness practice). The detection of cognitive effects associated with meditation may also depend on the specific processes tested, even within attention and executive control domains. These effects are likely to vary with task difficulty, the sensitivity of measures at detecting individual differences, and participants’ subjective experience; for example, how monotonous they find the task, with meditators possibly able to maintain attention better during boring tasks (Bailey et al., 2023a).

Identifying specific cognitive tasks that can differentiate meditators from non-meditators could significantly advance our understanding of the effects of meditation. Further, interventions that specifically target underlying mechanisms provide larger effects than those targeting outcomes, so a deeper understanding of mindfulness meditation mechanisms might facilitate the development of more effective interventions (Britton et al., 2018). Finally, since mindfulness meditation is now accepted as a form of mental training that results in improved attention and associated alterations in neural activity, characterisation of its underlying neurophysiological mechanisms could also be informative for our understanding of brain function more generally.

To investigate differences in cognition and brain activity associated with meditation, our previous research examined EEG activity during a Go/Nogo task (Bailey et al., 2019a). This task requires participants to respond to Go stimuli and inhibit responses to Nogo stimuli. The Go/Nogo task is known to elicit a pre-potent response tendency that must be suppressed in Nogo trials. The task engages sustained attention and executive function, particularly conflict monitoring and inhibition processes, with event-related potentials (ERPs) obtained from EEG recordings reflecting these processes (Bokura et al., 2001; R. J. Huster et al., 2013). Two ERPs are commonly examined in relation to the Go/Nogo task: the N2 and P3. The N2 ERP is a negative deflection prominent over fronto-central electrodes occurring approximately 200–300 ms post-stimulus. This response is larger in Nogo trials and is associated with conflict monitoring and unexpected stimuli processing (Donkers & Van Boxtel, 2004; Falkenstein, 2006; Smith et al., 2010). The P3 ERP is a positive deflection, which is present 300–600 ms after stimulus presentation and reflects attentional processes and expectation updating, with a more frontal distribution during cognitive and motor inhibition in Nogo trials (Datta et al., 2007; Kluger et al., 2020; Smith et al., 2008; Wickens et al., 1983).

In our previous study, mindfulness meditators showed higher overall accuracy at performing the Go/Nogo task compared to non-meditators (Bailey et al., 2019a). This is in alignment with other prior research examining Go/Nogo task performance in meditators (Andreu et al., 2019; Sahdra et al., 2011; Zanesco et al., 2013); but we note that improved Go/Nogo performance in meditators is not consistently reported (Korponay et al., 2019). In addition to these behavioural effects, we also observed three distinct differences in neural activity patterns between meditators and non-meditators. First, mindfulness meditators showed more negative voltages in occipital regions within 60 ms of stimuli presentation. Since visual information takes approximately 60 ms to reach the occipital cortex, we interpreted this altered distribution of neural activity as reflecting heightened anticipatory processes. Second, during the P3 window, meditators displayed equal amplitudes of their global neural response strengths between Go and Nogo trials, whereas non-meditators showed a larger global P3 amplitude for Go trials compared to Nogo trials. We interpreted this pattern as evidence that meditators exhibit reduced habitual response tendency in the task. Finally, meditators demonstrated a more frontally distributed P3 response for both Go and Nogo trials, which we interpreted as reflecting enhanced sustained attention processes. However, our previous study showed no meditation-related effects on the N2, despite the relationship between the N2 and conflict monitoring, and the suggestion that meditation enhances executive function (including conflict monitoring). One possible explanation for this null result is that our previous study used equal frequencies of Go and Nogo trials. Past research indicates that tasks with more equal Go and Nogo frequencies generate substantially less response inhibition and conflict monitoring demands than those with more frequent Go trials (Wessel, 2018). As such, we designed the present study to test whether a harder Go/Nogo task (with 25% Nogo trials) that imposes higher demands on conflict monitoring processes might be more sensitive in differentiating meditators and non-meditators based on N2 response amplitudes.

Additionally, since publication of our previous study, the predictive processing framework has gained prominence for offering a deep and parsimonious explanations for brain functions, with this framework increasingly applied to develop and test theoretical perspectives that attempt to explain the effects of meditation on the brain (Bailey, Hill, Godfrey, Perera, Hohwy, et al., 2024; Laukkonen & Slagter, 2021; Lutz et al., 2019; Manjaly & Iglesias, 2020). The predictive processing framework proposes that the brain functions by constructing a predictive model of its anticipated sensory experiences, including beliefs about how sensory experiences are altered based on one’s actions (Friston, 2010). Within this framework, the brain’s predictive model is continually updated by new sensory evidence, via the processing of mismatches between predictions and incoming sensory evidence (referred to as prediction errors) (Friston, 2010). The framework conceptualises the brain as a hierarchical system, where information that can update the predictive model is passed to higher levels of the neural hierarchy, weighted by the precision of both the prediction and the expected precision of sensory evidence (Wacongne et al., 2011). In the context of the predictive coding framework, the N2 and P3 can be seen as reflecting the processing of events that promote predictive model updates. The amplitude of ERPs may index the strength of predictive model updating as determined by the precision allocated to the processing of sensory inputs, while the frontal distribution of the ERPs might index how far up the cortical hierarchy the prediction error is processed.

Building on this background, the current study aimed to determine whether our previous findings concerning neural activity in the Go/Nogo task among mindfulness meditators could be replicated. Furthermore, we sought to examine whether a more challenging version of the Go/Nogo task (with less frequent Go trials) could elicit additional differences between meditators and non-meditators in the conflict monitoring and expectation related N2 component. We hypothesised that, compared to non-meditators, meditators would: 1) exhibit enhanced activity in occipital regions simultaneously with stimulus presentation, 2) display more frontally distributed P3 activity across all trials, 3) show less differentiation between Go and Nogo trials in global P3 amplitudes, and 4) differ from non-meditators in the contrast between easy and hard Nogo trials for the fronto-central N2 activity, reflecting altered processing of conflict monitoring demands. Finally, to enable interpretation of our findings, we applied exploratory analyses of the effect of different Go/Nogo task parameters on group differences in neural activity with the aim to better understand the functional implications of the neural activities that showed group differences.

## Methods

### Participants

Thirty-nine experienced mindfulness meditators and 37 non-meditators were recruited through community advertising at meditation organisations, universities, and through social media. From this cohort, a total of 64 individuals (35 meditators and 29 non-meditators) provided enough artifact free EEG activity for analysis and met the study inclusion criteria (more details are provided in the EEG processing section of the methods). This participant sample partially overlaps with the sample reported in four of our other previously published studies that examined different cognitive tasks, but differs due to study-specific exclusions (Bailey et al., 2023a; Bailey et al., 2023e; Bailey, Hill, Godfrey, Perera, Hohwy, et al., 2024; Payne et al., 2020).

Notably, an analysis of cortical travelling waves has already been reported from a combination the Go/Nogo data reported in this study and the Go/Nogo data reported in our previous study (Bailey, Hill, Godfrey, Perera, Hohwy, et al., 2024). Approximately one-third of the participants also contributed to another published study focused on resting-state EEG data (Bailey et al., 2023d). The data were collected by students who were trained in EEG and supervised by experienced investigators.

To meet inclusion criteria, meditators required a current meditation practice of at least 2 hours of meditation per week for the most recent three-month period (final sample mean = 7.124 hours per week, SD = 5.265). All participants reported a meditation practice duration of at least 18 months (final sample mean = 8.171 years, SD = 8.331). Meditation practices were required to be mindfulness-based, adhering to Kabat-Zinn’s (1994) definition: “paying attention in a particular way: on purpose, in the present moment, and nonjudgmentally”. We additionally required that the meditation practice was focused on sensations of the breath or body. Trained meditation researchers screened and interviewed participants to ensure meditation practices adhered to these criteria. Participants were included in the non-meditator group if they had less than two hours of lifetime meditation experience. Any uncertainties as to whether a participant’s mindfulness practice met the definition used in the study were resolved through discussion and consensus between the principal researcher and one other researcher. A conservative approach was adopted, meaning that if uncertainties persisted after discussion, the individual was excluded from the study.

Exclusion criteria included any previous or current neurological or mental illness, current use of psychoactive medication, or psychoactive drug use within the past two weeks. All participants were interviewed with the Mini-International Neuropsychiatric Interview (MINI) (Sheehan et al., 1998) and were excluded if they met the criteria for any psychiatric illness. Since anxiety and depression have been shown to affect brain activity, the Beck Anxiety Inventory (BAI) and Beck Depression Inventory II (BDI-II) were also administered (Beck et al., 1996; Steer & Beck, 1997) and participants who scored above the moderate range in the BAI (>25) or above the mild range in the BDI-II (>19) were excluded. All participants were aged between 20 and 64 years of age. Ethical approval for the study was provided by the Ethics Committees of Monash University and the Alfred Hospital. All participants provided written informed consent prior to participation in the study.

Prior to EEG recording and task completion, participants provided demographic information and completed the self-report questionnaires. This included providing their age, gender, years of education and meditation experience (total years of practice, frequency of practice, and amount of time spent practicing per week). In addition to the BAI and BDI-II, participants completed the Five Facet Mindfulness Questionnaire (FFMQ) (Baer et al., 2006). Means and standard deviations for these measures are provided in Table 1. Groups did not significantly differ in term of age (*T_y_* = 1.898, *df* = 29.260, *p* = 0.068), or years of education (*T_y_* = 1.536, *df* = 37.880, *p* = 0.133). However, while the difference was not significant, the mean age did vary considerably between the groups (meditator mean = 37.230, SD = 15.000, non-meditator mean = 29.900, SD = 11.130). As such, in addition to our primary analysis (which including all eligible participants), we performed confirmatory analysis where each participant was randomly matched in age (to within five years) against a single participant from the other group. This process excluded one non-meditator and seven meditators, resulting in 28 participants in each group and matched ages between the groups (meditator mean = 31.893, SD = 11.487, non-meditator mean = 30.143, SD = 11.250).

**Table 1.** Self-report data from the sample of 35 meditators and 29 non-meditators who were included in the study.

|  | Meditators: Mean (SD) | Non-meditators: Mean (SD) | Statistics |
| --- | --- | --- | --- |
| Age | 37.229 (14.994) | 29.897 (11.127) | $T_y = 1.898, p = 0.067$ |
| Gender (F/M) | 12/17 | 11/24 | $ChiSq = 0.682, p = 0.409$ |
| Years of education | 15.971 (3.044) | 17.259 (2.348) | $T_y = 1.536, p = 0.1328$ |
| Years of meditation Practice | 8.171 (8.331) | 0 |  |
| Hours of meditation practice per week | 7.124 (5.265) | 0 |  |
| FFMQ | 148.886 (29.462) | 131.552 (15.715) | $T_y = 4.854, p < 0.001$ |
| BDI-II | 2.886 (3.756) | 4.552 (4.918) | $T_y = 0.667, p = 0.509$ |
| BAI | 5.686 (6.178) | 5.786 (5.527) | $T_y = 1.338, p = 0.194$ |

### Procedure

#### EEG Data Acquisition

Sixty-channel EEG (Neuroscan, Ag/AgCl Quick-Cap) was recorded using a standard 10-10 electrode montage while participants completed an easy and a hard version of a Go/Nogo task. EEG was recorded via a SynAmps 2 amplifier (Compumedics, Melbourne, Australia). Electrodes were referenced online to a midline electrode between Cz and CPz. Impedances below 5kΩ were obtained prior to the start of recording. EEG data were sampled at 1000 Hz with online bandpass filters from 0.1 to 100 Hz.

### Go/Nogo Task Administration

Both the easy and hard Go/Nogo tasks used simplified emotional faces as stimuli (see Figure 1), identical to those used in our previous study (Bailey et al., 2019a). However, while our previous study presented only an easy version of the task with a 50% Nogo frequency, the current study included both the easy version and a harder version with a 25% Nogo frequency. Participants completed four separate blocks of the tasks. The first two and the last two blocks were randomly counterbalanced, so that one block required participants to respond to the happy faces and withhold responses to the sad faces, while the other block required participants to respond to the sad faces and withhold responses to the happy faces. The first two blocks presented the easy version of the task, each showing a total of 50 happy and 50 sad faces (for a total of 100 Go and Nogo trials) in replication of our previous study. Following this easy version of the task, participants completed the harder version of the task in two separate blocks. These hard task blocks presented 50 Nogo trials and 150 Go trials (for a total of 100 Nogo and 300 Go trials). Each stimulus was presented for 250 ms with an inter-trial interval of 900 ms (with a 50 ms jitter to reduce the effects of entrainment). Our previous research showed no interactions between group and the emotion used as stimuli, so Go and Nogo data were separately averaged across happy and sad stimuli to reduce the complexity of analyses (Bailey et al., 2015; Bailey et al., 2019a; Bailey et al., 2014).

**Figure 1.**
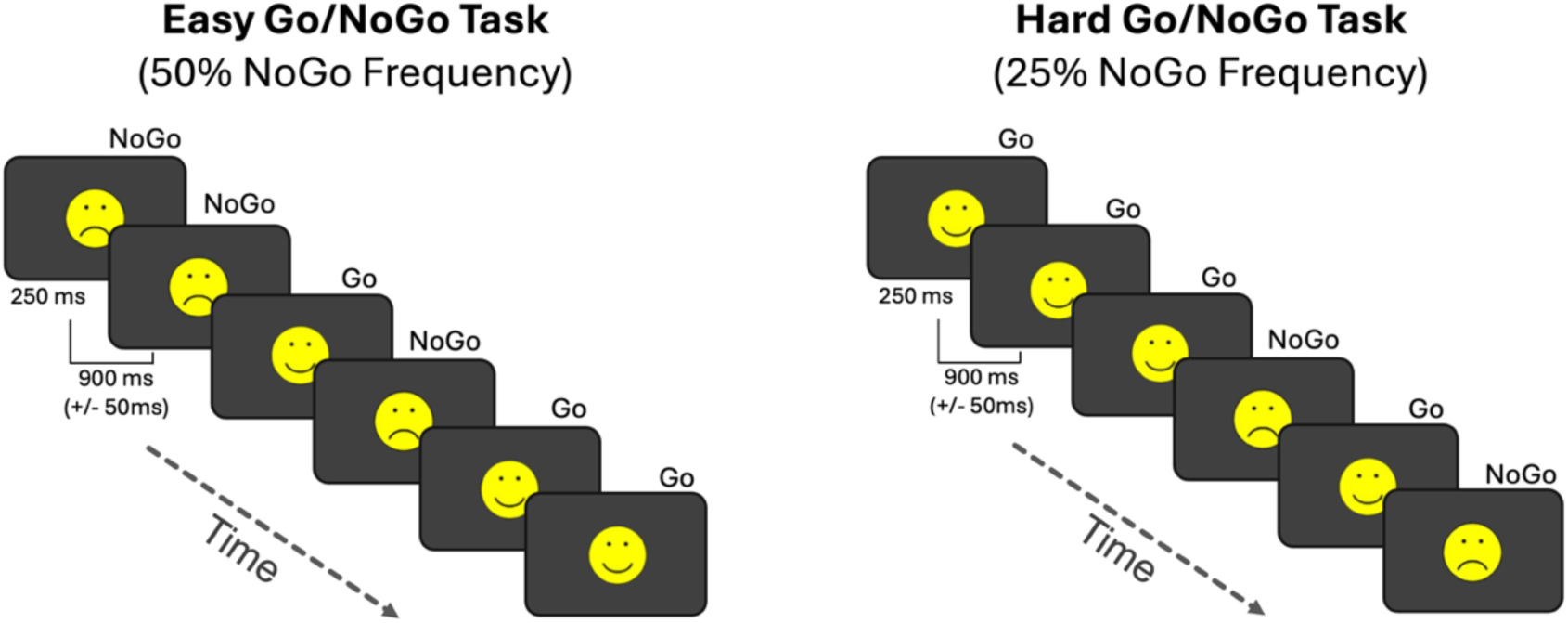
Task design. Go and Nogo stimuli were randomly presented, each for 250ms with an inter-trial interval of 900ms with a 50ms jitter. The easy version of the task presented the Go and Nogo trials with equal frequency, while the hard version of the task only presented 25% of the trials as Nogo stimuli.

#### EEG Data Pre-Processing

EEG data were pre-processed using the RELAX pipeline (Bailey et al., 2023b), a toolbox that uses EEGLAB functions (Delorme & Makeig, 2004), with selected application of Fieldtrip functions (Oostenveld et al., 2011). Data were pre-processed using the *wICA_ICLabel* default RELAX settings for ERP analysis, which have been demonstrated to provide effective reduction of artifact and preservation of signal (Bailey et al., 2023b), and the largest effect sizes for between condition comparisons of Go and Nogo conditions in a Go/Nogo task (Bailey et al., 2023c). For the sake of brevity, the full EEG pre-processing details are reported in the supplementary materials.

Following cleaning, the continuous data were epoched from -100 to 800 ms surrounding each stimulus around correct and incorrect Go and Nogo responses from the easy and hard version of the task separately. Epochs with a voltage exceeding +/- 100 μV at any electrode were rejected, as were epochs containing improbable voltage distributions or kurtosis values > 5SD from the mean in any single electrode or more than 3SD from the mean over all electrodes (Bailey et al., 2023c). A total of 12 participants were excluded from analysis at this stage due to not meeting data integrity criteria (four meditators and eight non-meditators). Six non-meditators and two meditators were excluded due to technological faults during recording which left their EEG data unusable. One non-meditator was excluded for revealing a previous history of meditation practice after the EEG recording. One non-meditator was excluded due to high anxiety scores in the BAI. One meditator was excluded for revealing a previous history of neurological illness after the EEG recording. One meditator was excluded for continuing to respond to the happy faces after the stimulus-response pairing was switched. All remaining participants provided more than 50 correct and artifact free EEG trials in each condition for analysis.

Next, the cleaned and epoched EEG data were baseline corrected by using a regression baseline correction method, which controls for the influence of the average of the −100 to 0ms period on the period of interest in each epoch (time-locked to the response) using the RELAX function “RELAX_RegressionBL_Correction” for each electrode and each participant separately. Within the regression, the condition of each epoch (Go or Nogo stimuli) was set as the first factor, and the easy or hard task was set as the second factor, with a constant included in the regression model (but not rejected) to control for potential voltage drift but still preserve experimental effects (Bailey et al., 2023c; Bailey et al., 2023e). This approach removes the influence of the baseline period from the neural activity of interest but does not introduce differences from the baseline into that activity, in contrast to the subtraction baseline correction method, which transposes baseline differences into the active period of interest (Alday, 2019; Bailey et al., 2023e). A more detailed explanation of the regression baseline correction method is provided in the supplementary materials.

### Data analysis

Demographic variables were tested to ensure groups were matched (but differed in trait mindfulness as is expected) using Yuen’s test (*T_y_*) in R with the WRS2 package (Mair & Wilcox, 2020). Yuen’s test is equivalent to Welch’s t-test but is implemented with trimmed means to control for violations in the assumptions of the Welch’s t-test, an approach recommended as the default for psychological research (Mair & Wilcox, 2020). Behavioural data were similarly tested using trimmed mean approaches with the WRS2 package, with the bwtrim test to test for group main effects and interactions between the easy and hard task in repeated measures ANOVA designs (Mair & Wilcox, 2020). We provide the test statistic for the results of the bwtrim tests as *bwtrim.* Comparisons between mean reaction times to Go trials in the easy and hard version of the task were also tested with the bwtrim test. Accuracy was measured using the loglinear adjusted *d*-prime (to control for perfect performance in either the true hit or false alarm rate (Stanislaw & Todorov, 1999) and compared between groups and the easy and hard version of the task using the bwtrim test. Three-way interactions were tested using traditional repeated measures ANOVAs in JASP (Love et al., 2019), since bwtrim does not allow for designs that involve more than one repeated measures factor.

ERP comparisons were conducted using the randomization graphical user interface (RAGU) (Koenig et al., 2011). RAGU is a rank order randomisation statistics toolbox designed to analyse EEG data, enabling t-test or ANOVA statistical designs that include all electrodes and timepoints within an ERP epoch in the analysis, while robustly controlling for multiple comparisons (Koenig et al., 2011). RAGU uses scalp field differences (measured via the global field potential; GFP) across all electrodes to reduce the number of comparisons in the electrode space. It conducts statistical tests at each timepoint in the epoch followed by a control for multiple comparisons using a global duration control, which tests for the proportion of real data that shows significant effects that last longer than 95% of the significant effects in the randomised data (Koenig et al., 2011). This allows for analyses that include all electrodes and timepoints in the ERP epoch without being limited by researcher assumptions about which time windows or electrodes might show significant effects (Koenig et al., 2011). RAGU also allows independent comparisons of potential differences in the strength of the global neural response to stimuli (using the GFP test), as well as potential differences in the distribution of neural activity across the scalp after controlling for the potential influence of variability in global neural response strengths (using the L2 normalised topographical analysis of variance [TANOVA] test) (Koenig et al., 2011). The GFP is equivalent to the standard deviation across all electrodes, reflecting the overall voltage amplitude, with larger values indicating a stronger neural response (Koenig et al., 2011). A full explanation of how RAGU performs these statistical tests is provided in the supplementary materials.

The *p*-values from our primary hypotheses (with data averaged within a priori hypothesised time windows of interest) were submitted to False Discovery Rate (FDR) multiple comparison controls (Benjamini & Hochberg, 2000) to control for experiment-wise multiple comparisons (referred to as *p*_FDR_). Following initial statistical tests using RAGU, we performed generalised eigenvector decompositions (GED) to spatially filter data separately for each individual, enabling us to extract a single time-series for each epoch representing the source space phasic ERP signals that most strongly represented the topographical patterns shown by RAGU to differ between the groups (Cohen, 2022). We then used these GED decomposed time-series to perform single trial analyses with linear mixed models (LMM) and generalised linear mixed models (GLMM), enabling us to interpret the functional relevance of the patterns that differed between the groups. However, while the N2 effects involving group provided a clear scalp map target for GED extraction, the effects for the P3 difference did not. As such, single electrode ERP amplitudes averaged within the significant effect period were used in the linear mixed models instead, in addition to an extraction of a measure of the topographical centroid of the P3 activity. For brevity, we report the full methods and results of these exploratory analyses in the supplementary materials and provide only brief summaries of the results that are important for interpretation of our primary analyses in the results section of our main manuscript.

## Results

### Meditators Performed Better in the Easy Go/Nogo Task When Groups Were Matched for Age

There was no significant difference between the meditator and non-meditator groups in task accuracy when analysing *d*-prime scores (*bwtrim* = 2.609 [*df* = 1,35.088], *p* = 0.115, see Figure 2 and Table 2), nor was there a significant interaction between group and task difficulty (*bwtrim* = 2.925 [*df* = 1,35.686], *p* = 0.096).

**Figure 2.**
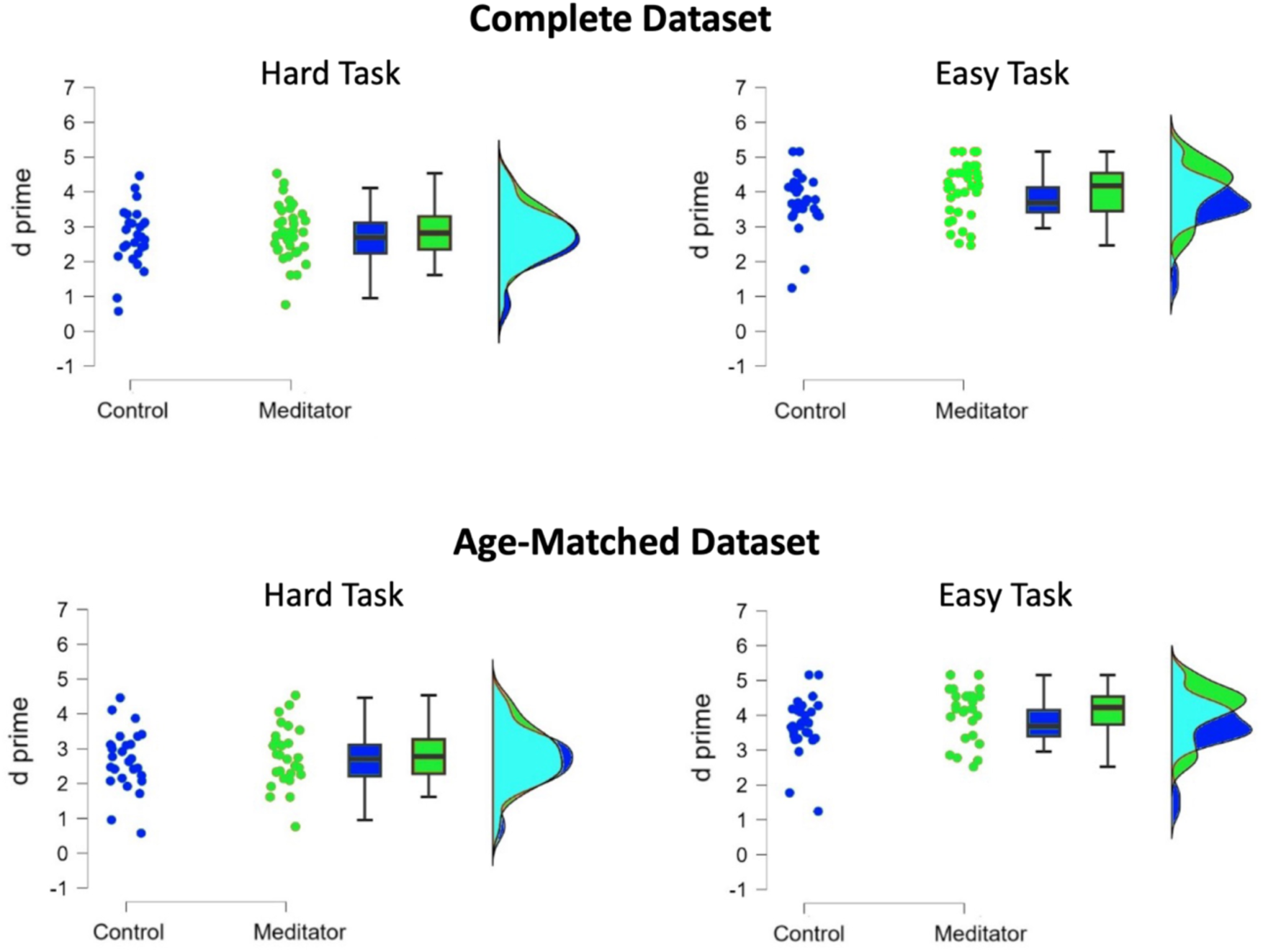
Task performance (*d*-prime) for the easy and hard tasks for each group across all participants (top) and within the age-matched groups only (bottom). No significant main effect or interaction involving group was present for the complete dataset (both *p* > 0.05). However, when analyses were restricted to age-matched groups, a significant interaction was observed between group and task difficulty (*bwtrim* = 4.841 [*df* = 1,33.357], *p* = 0.035), with meditators showing higher *d*-prime scores in the easy task than non-meditators.

**Table 2.**
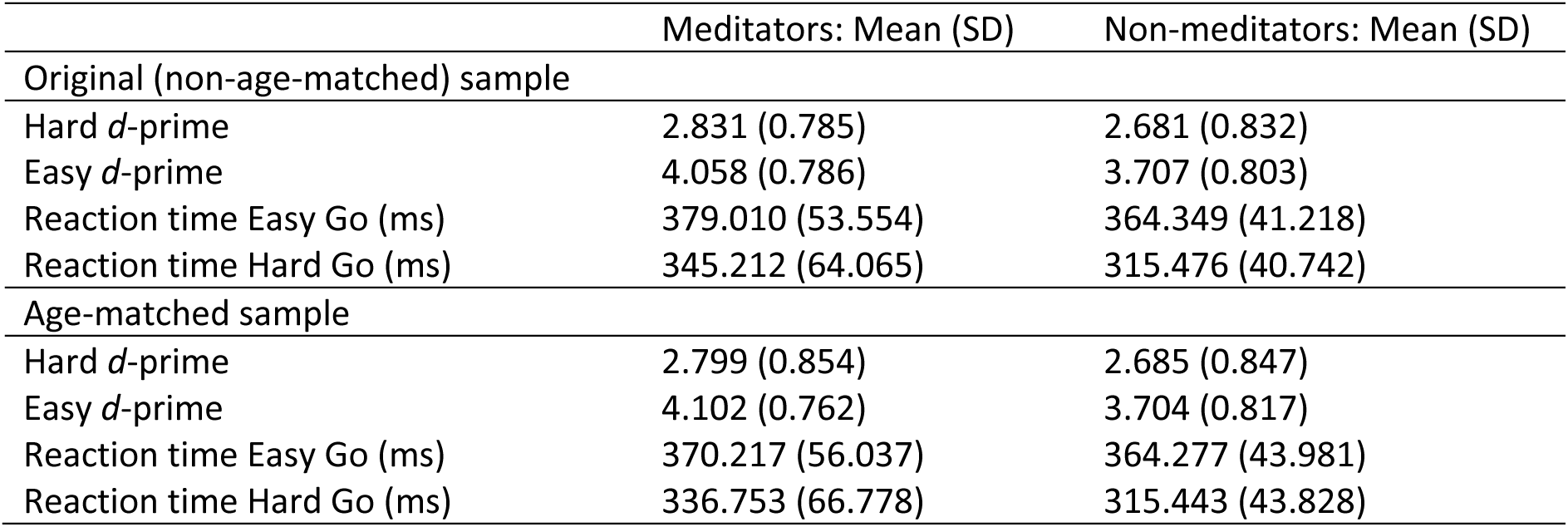
Behavioural performance from each group for each of the tasks.

|  | Meditators: Mean (SD) | Non-meditators: Mean (SD) |
| --- | --- | --- |
| Original (non-age-matched) sample |  |  |
| Hard *d*-prime | 2.831 (0.785) | 2.681 (0.832) |
| Easy *d*-prime | 4.058 (0.786) | 3.707 (0.803) |
| Reaction time Easy Go (ms) | 379.010 (53.554) | 364.349 (41.218) |
| Reaction time Hard Go (ms) | 345.212 (64.065) | 315.476 (40.742) |
| Age-matched sample |  |  |
| Hard *d*-prime | 2.799 (0.854) | 2.685 (0.847) |
| Easy *d*-prime | 4.102 (0.762) | 3.704 (0.817) |
| Reaction time Easy Go (ms) | 370.217 (56.037) | 364.277 (43.981) |
| Reaction time Hard Go (ms) | 336.753 (66.778) | 315.443 (43.828) |

However, the main effect of task difficulty was significant, with lower *d*-prime scores for the harder task, as expected (*bwtrim* = 206.767 [*df* = 1,35.686], *p* < 0.001). When analyses were restricted to age-matched groups, there was still no main effect of group (*bwtrim* = 2.017 [*df* = 1,31.532], *p* = 0.165). However, within the age-matched groups, there was a significant interaction between group and task difficulty (*bwtrim* = 4.841 [*df* = 1,33.357], *p* = 0.035), with meditators showing higher *d*-prime scores in the easy task compared to non-meditators. This finding contradicts our expectations that the hard task would place higher cognitive demands on participants and reveal larger effects. However, it does replicate our previous research, which demonstrated a between-group difference in performance in the easy Go/Nogo task (Bailey et al., 2019a). Within our analysis of all participants, no significant between-group difference was detected in reaction time (*bwtrim* = 1.791 [*df* = 1,33.981], *p* = 0.190), nor was there an interaction between group and task difficulty (*bwtrim* = 1.765 [*df* = 1,31.891], *p* = 0.193). The main effect of task difficulty was significant, as expected (*bwtrim* = 58.446 [*df* = 1,31.891], *p* < 0.001), with reaction times slightly faster in the hard version of the task. When analyses were restricted to age-matched groups, there were still no significant main effects or interactions involving group (all *p* > 0.05).

#### Meditators Showed More Negative Fronto-Central N2 ERP Topographies in Hard Nogo and Easy Go Trials

Our analysis of the fronto-central N2 ERP indicated that meditators demonstrated a more negative fronto-midline N2 activity in response to easy Go trials and hard Nogo trials (Figure 3). In particular, a significant interaction between Group, Go/Nogo trial type, and difficulty was present in the TANOVA from 198 to 241ms (*p-*value averaged across period of significance = 0.001, *p*_FDR_ = 0.011, *np^2^* = 0.111, global duration control = 38ms). This aligns with the N2 window, with activity across this period showing the typical negative fronto-central N2 topographical distribution. We explored the drivers of this interaction with post-hoc analyses using *t*-test designs in RAGU to restrict comparisons to two relevant conditions at a time. These indicated that the interaction was driven by between group differences in both the hard Nogo trials (*p* = 0.013, *np^2^* = 0.041) and the easy task Go trials (*p* = 0.005, *np^2^* = 0.055), with meditators demonstrating more negative fronto-midline N2 activity in response to both trial types. There were no other significant main effects of group nor interactions between group and Go/Nogo condition that exceeded the global duration control threshold at any point during the epoch for the GFP or TANOVA tests (Figure 3).

**Figure 3.**
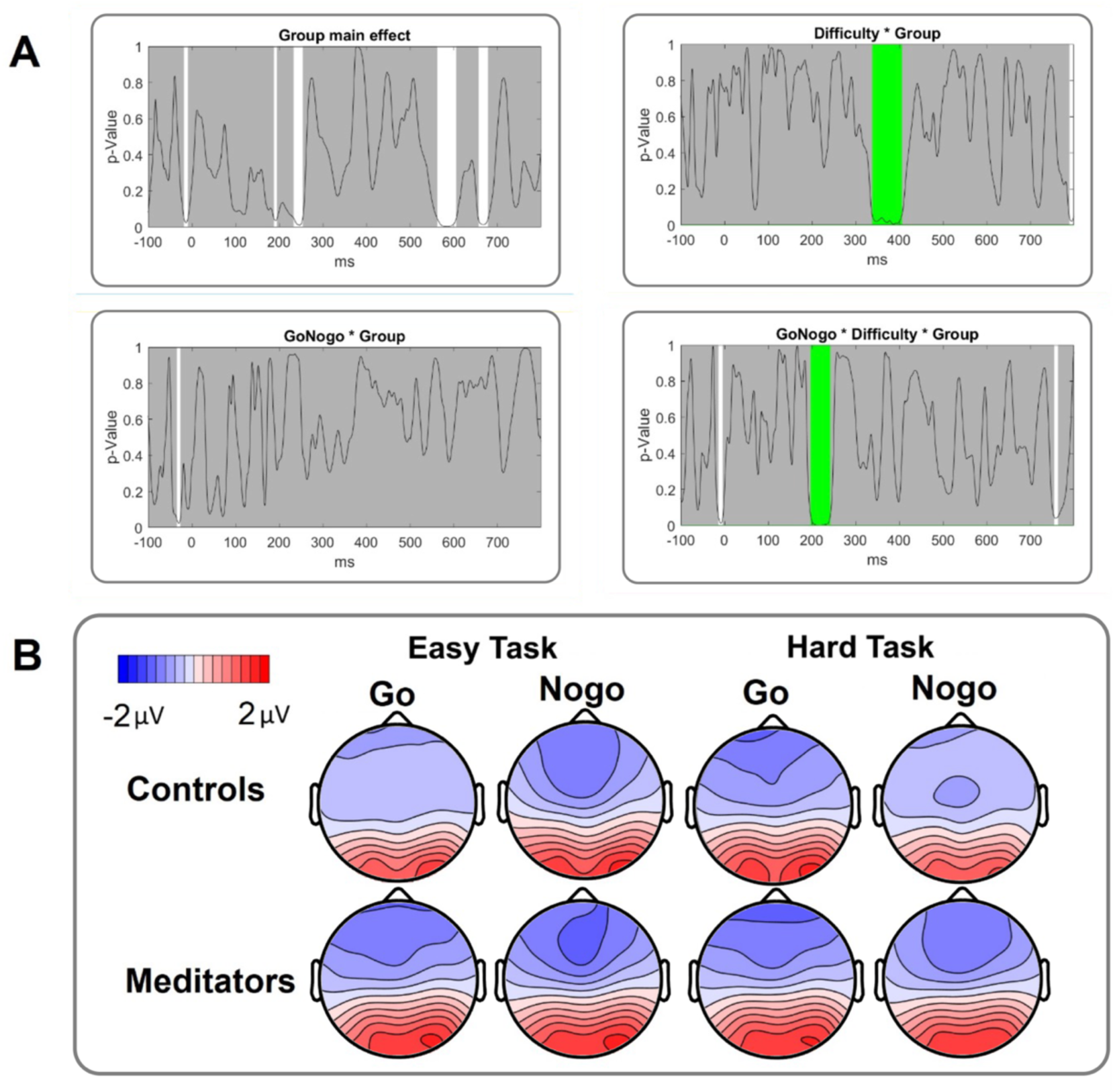
A) TANOVA *p*-graphs for each of the main effects of group and interactions between conditions and group. The black line reflects the *p*-value at each timepoint in the epoch, white patches reflect periods where the *p*-value is below 0.05, while green patches reflect periods where *p*-values fall below 0.05 for longer than the global duration control (which controls for multiple comparisons in the time domain). B) Topographic plots of the N2 ERP response for each group and trial type, averaged within the significant 198 to 241ms window. Post-hoc tests indicated the significant interaction in the N2 window was driven by between group differences in both the hard Nogo trials (*p* = 0.013, *np^2^* = 0.041) and the easy task Go trials (*p* = 0.005, *np^2^* = 0.055), with meditators showing more negative fronto-midline N2 activity in response to both of these trial types.

Within analysis of the age-matched groups, comparisons of averaged data within the N2 period (198 to 241ms) revealed a significant main effect of group (*p* = 0.017, *np^2^*= 0.048), and the same three-way interaction between group, Go/Nogo condition and difficulty as the non-age-matched groups (*p* = 0.011, *np^2^* = 0.086). These results suggest the meditators showed a stronger topographical concentration of the fronto-central negative N2, with this pattern being most pronounced in the easy Go and hard Nogo trials. Although the N2 response is associated with conflict monitoring, the between group difference observed in Go trials during the task suggests it may be driven by an attention-related mechanism rather than a difference in conflict monitoring processes. This possibility is further explored in the single-trial analyses reported below.

### Meditators Displayed a More Frontally Distributed P3 ERP Response in the Easy Task Compared to the Hard Task, Whereas Non-Meditators Showed the Opposite Pattern

In addition to the effect in the N2 window, our analyses indicated topological differences in the P3 ERP distribution between the easy and hard Go/Nogo tasks across the meditator and non-meditator groups. In particular, meditators showed a more posteriorly distributed P3 during the hard task than the easy task, while non-meditators showed a more posteriorly distributed P3 in the easy task than the hard task (Figure 4). Specifically, the TANOVA showed a significant interaction between group and task difficulty within the P3 time window (339 to 405 ms; Figure 4), an effect that lasted longer than the global duration controls (*p*-value averaged across period of significance = 0.004, *p*_FDR_ = 0.015, *np^2^* = 0.079, global duration control = 34ms). Post-hoc analysis revealed group differences in the topographical pattern of the between-task effect, with non-meditators showing a significant difference between the easy and hard tasks (*p* = 0.002, *np^2^* = 0.265). Specifically, their P3 in the hard task exhibited more positive voltages over motor regions (with a minimum in the difference map at electrode C6) and less positive voltages at parieto-occipital electrodes (with a maximum in the difference map at electrode PO4) compared to the easy task (indicating a more posteriorly distributed P3 in the easy task). In contrast, meditators displayed a similar overall P3 topographical pattern to non-meditators, but a weaker effect between easy and hard trials (*p* = 0.002, *np^2^* = 0.136). Their P3 difference map showed less negative voltages in prefrontal electrodes during the easy task (with a maximum in the difference map at AF3, indicating a more frontally distributed P3 in the easy task) and a similar pattern to non-meditators over motor regions (with a minimum in the difference map at C3). ERP graphs for each of the electrodes involved in this effect across each task can be viewed in the supplementary materials (Figure S2). Our analysis of the same interaction between group and task difficulty averaged within the P3 window within the age-matched groups showed the same interaction effect (*p* = 0.004, *np^2^* = 0.094). Our single trial analysis reported below attempts to shed light on the functional implications of the between group difference in the P3 distribution depending on the task difficulty.

**Figure 4.**
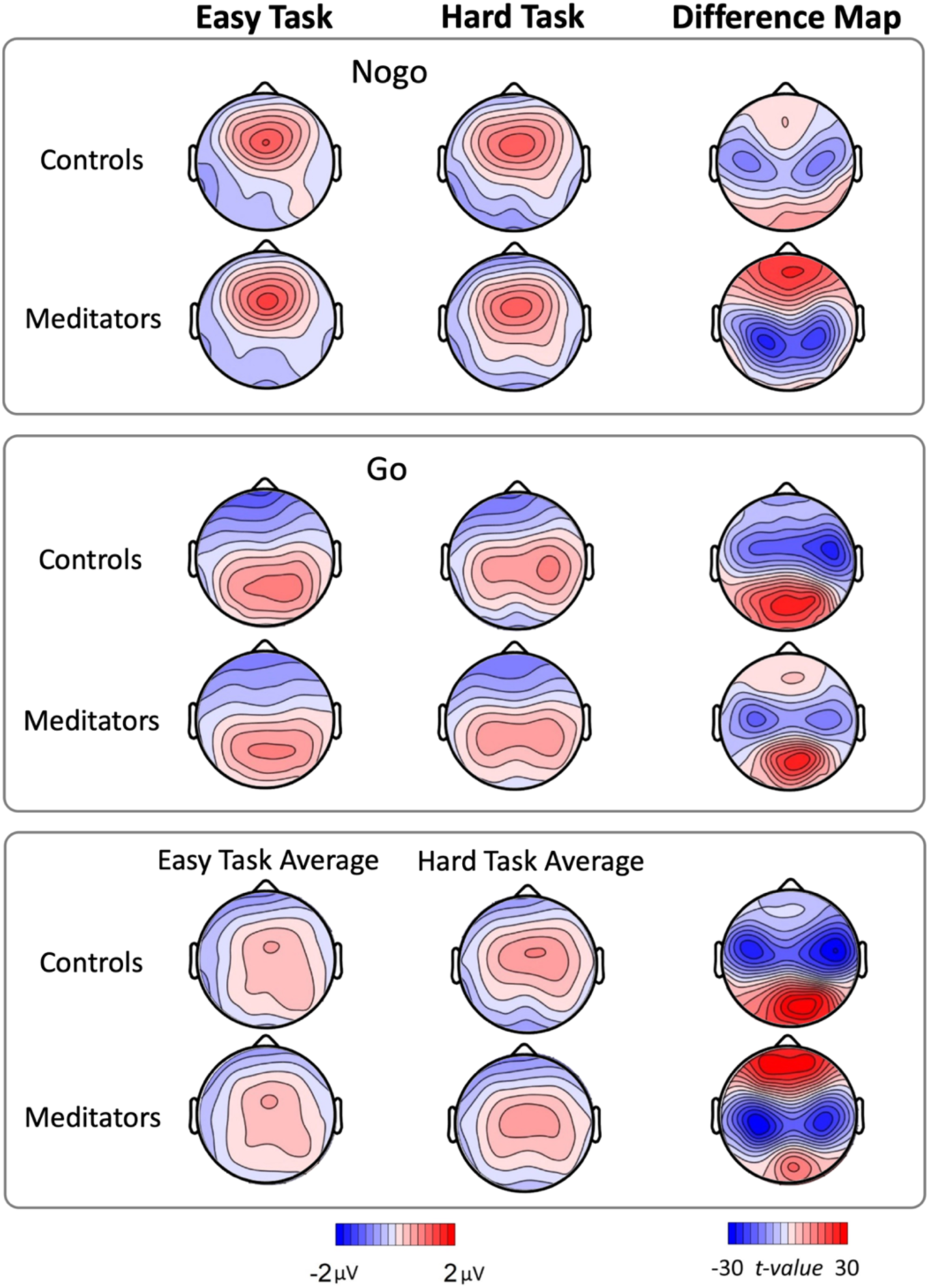
P3 topography averaged within the 338 to 406ms window for each condition, with difference maps reflecting activity in the easy task minus activity in the hard task. There was a significant interaction between group and task difficulty in the topographical distribution of the P3 (*p*_FDR_ = 0.015, *np^2^* = 0.079), where meditators showed more frontally distributed P3 topographies in the easy version of the task.

### Single-Trial Analysis Suggests Meditators’ Increased Hard Nogo Fronto-Central N2 is Related to Increased Attention to Stimuli

Our exploratory single trial examination of the GED N2 ERP using LMM and GLMM analyses revealed significant relationships between the GED N2 and task parameters, as well as interactions between group and task parameters (p < 0.05). Full results are reported in the supplementary materials and are only summarised here for brevity. First, the GED N2 ERP was significantly larger (more negative) in meditators than non-meditators across both Go and Nogo trials, in alignment with the main effect of group detected in our RAGU analyses of the age-matched dataset (Figures 5 and 6).

**Figure 5.**
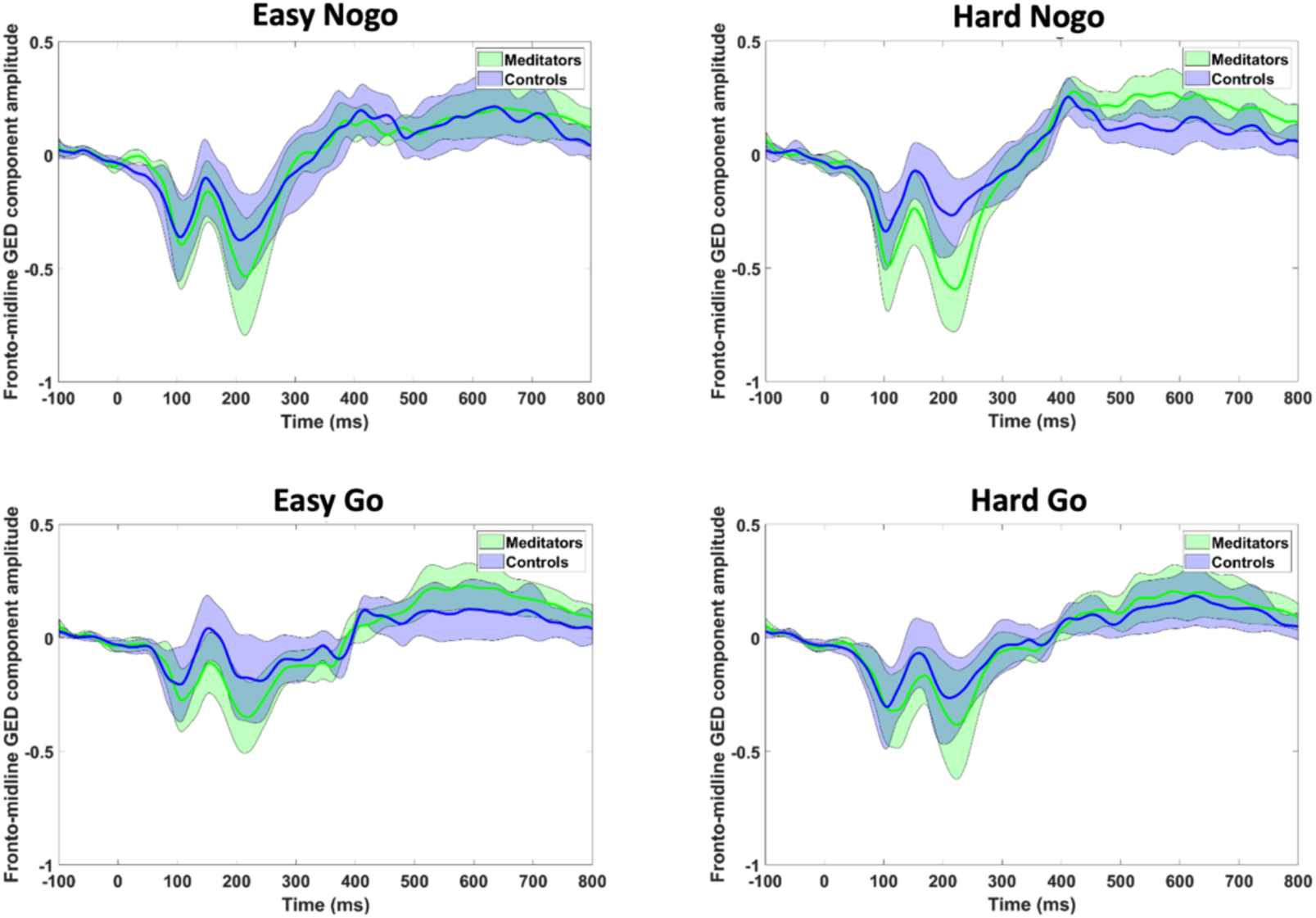
Generalised eigenvector decomposition extracted fronto-midline N2 event-related potential amplitudes for each condition across both groups. Error shading reflects 95% confidence intervals.

**Figure 6.**
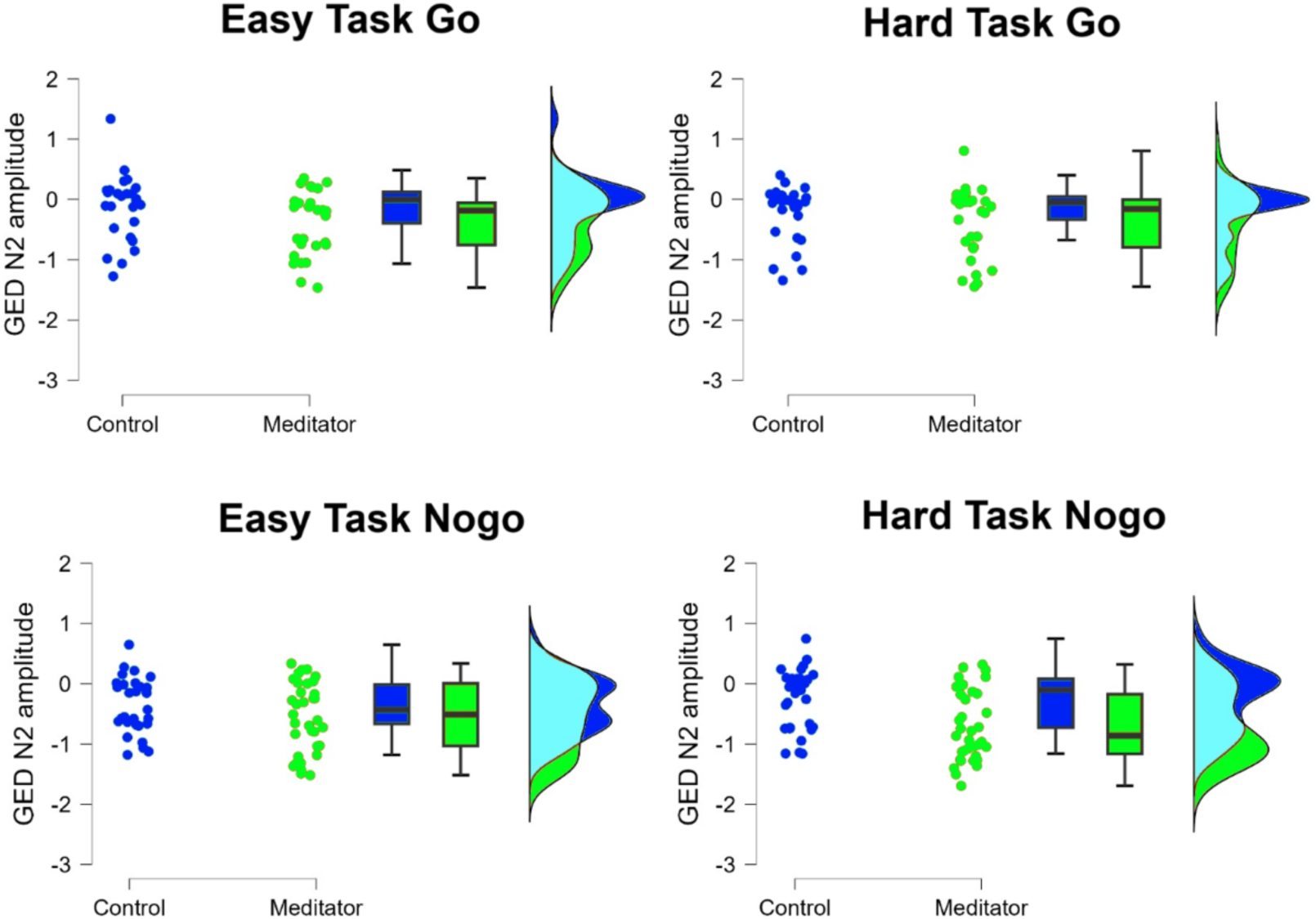
Generalised eigenvector decomposition extracted fronto-midline N2 amplitudes averaged across the 150 to 250ms window for hard Nogo trials and easy Nogo trials. When task difficulty and group were modelled alongside number of prior Go trials in our LMM analysis, results showed a main effect of group where the N2 component was significantly larger (more negative) for meditators than controls (F(1,62) = 6.59, p = 0.013). See supplementary materials for further details.

Second, there was a trend level interaction between group and task (p = 0.051), with a pattern indicating meditators had more negative Nogo GED N2 responses in the hard task compared to the easy task, while non-meditators showed more negative Nogo GED N2 responses in the easy task compared to the hard task (Figure 6). While not significant, this pattern nevertheless aligns with the group-by-task-by-trial type interaction demonstrated in our RAGU analyses. We note that the shift from the significant interaction in the averaged ERP analysis to a trend in the single trial LMM analysis is likely due to the proportion of the variance explained by the number of prior Go trials in the LMM, which the averaged ERP analysis did not examine.

Third, the GED N2 was larger in amplitude in hard Nogo trials than easy Nogo trials, and in Nogo compared to Go trials, consistent with the perspective that the N2 reflects conflict monitoring demands. However, the trial-level amplitude of the GED N2 differed in the easy and hard tasks depending on prior trial sequence. In the hard version of the task, the N2 amplitude diminished as the number of prior consecutive Go trials increased up to six trials, stabilising (or slightly increasing again) thereafter; in the easy task, the N2 amplitude increased for up to two consecutive preceding Go trials, returning to original levels thereafter.

Similar patterns were observed across both the Go and Nogo GED N2, although the effect was small in amplitude. This suggests that, beyond conflict monitoring demands, the N2 was modulated by interactions between local and global stimuli frequencies.

Fourth, larger Nogo GED N2 amplitudes were associated with correct responses in the hard task only. Interestingly, there was no interaction between the number of prior Go trials, Nogo GED N2 amplitudes, and correct responses, suggesting that increased expectations for Go trials did not increase the likelihood that larger amplitude Nogo GED N2s were required for a correct response. This suggests the relationship between larger N2 amplitudes and correct responses in the hard task was driven by global Go and Nogo frequencies (where the hard task presented 25% Nogo trials overall) rather than by local frequencies (since an increasing number of preceding Go trials did not alter the relationship between Nogo N2 amplitudes and correct non-responses).

### Single-Trial Analysis Indicates That Meditators’ More Frontally Distributed P3 During the Easy Task is Associated with Enhanced Attention

The full results for our LMM and GLMM analyses of single trial P3 activity are reported in our supplementary materials, and we report only a summary here for brevity. Given the interactions between group and task difficulty for the topographical distribution of the P3 ERP detected by our primary analyses, we focused our single trial LMM and GLMM analyses on the topographical P3 centroid (which indicates the degree of the frontal/posterior distribution of the P3). In alignment with our primary RAGU analyses, analysis of the Nogo P3 centroid showed an interaction between group and task difficulty, with meditators showing a more frontally distributed Nogo P3 in the easy task compared to the hard task, while non-meditators showed a more frontally distributed Nogo P3 in the hard task compared to the easy task (p < 0.05).

It is worth noting here that our easy task contained a higher proportion of Nogo trials compared to the hard task, which likely reduced participant expectations for Go trials. For non-meditators, the more frontal Nogo P3 distribution in the hard task aligns with the predictive processing framework, where prediction errors propagate further up the cortical hierarchy due to stronger top-down expectations, which are driven by higher stimuli frequencies. In contrast, however, the finding that meditators have more frontal P3 responses in the easy compared to the hard task conflicts with these predictions and warrants further explanation. The following results report significant effects (p < 0.05) from our single trial analysis of the P3 centroid, undertaken to explore the potential functional relevance of variations in the frontal distribution of the P3 to help with the interpretation of our results.

First, the Nogo P3 centroids showed a more frontal distribution than Go P3 centroids across both groups. This finding suggests that, given participants’ response tendencies or expectations/predictions for Go trials, prediction errors for Nogo stimuli were passed to higher levels in the cortical hierarchy. However, hard task Go P3 centroids were also more frontally distributed than easy Go P3 centroids across both groups, and both Go and Nogo P3 centroids shifted more frontally with an increasing number of prior Go trials. This result is more complex to interpret, as it suggests that in situations where participants would be hypothesised to have stronger expectations for Go trials (in the hard task and after more repeated Go trial presentations), prediction errors for both Go and Nogo trials were passed to higher levels in the cortical hierarchy compared to the situations where participants would be hypothesised to have weaker expectations for Go trials (in the easy task and with fewer prior Go trial presentations). Interestingly, the frontal shift in the P3 centroid with increasing repeats of prior Go trials was stronger for the easy task than the hard task, suggesting that prediction errors were processed higher in the cortical hierarchy when the local stimulus probabilities preceding the current trial were less likely based on global stimulus probabilities (since the easy task presented 50% Go trials, higher numbers of repeated Go trials in the easy task were less likely than in the hard task). Finally, more frontal Nogo P3 centroids were associated with a higher proportion of correct responses, and this effect was larger in the easy task than the hard task.

### Replication Testing

Finally, it is worth noting that the current results differed from our previous results (Bailey et al., 2019a). Our previous study only included the easy version of the Go/Nogo task, so could not have detected the significant results found in our current data (which all involved interaction effects between group and task difficulty). However, our previous study did report a very early difference between meditators and non-meditators in the topographical distribution of activity (-1 to 61ms around the stimuli), as well as a more frontal P3 distribution across both Go and Nogo trials (from 416 to 512ms), and more equal P3 global amplitudes between Go and Nogo stimuli (from 336 to 449ms), which could have been detected in the easy task within our current dataset. To test potential explanations for this discrepancy, we conducted exploratory tests, applying pre-processing parameters to the current data that were more similar to those used in our previous study. The results of these analyses are reported in full in the supplementary materials and summarised here for brevity. First, our exploratory analysis indicated that when a subtraction baseline correction method was applied to the data from the current study, the difference in the distribution of activity in occipital electrodes from -1 to 61ms replicated in the current dataset (*p* = 0.004, *np^2^* = 0.065).

Since the subtraction baseline method transposes a mirror image of the topography from the baseline correction period into the active period (Alday, 2019; Bailey et al., 2023c; Bailey et al., 2023e), we suspect that instead of representing a true effect at -1 to 61ms post stimulus, the effect is driven by a difference between the groups in the interaction between activity at the end of the previous trial (during the baseline correction period) and the activity immediately after stimulus presentation. These differences may reflect effects in the late positive potential/slow wave activity, where a nearly significant effect was present in our analysis of the current data using a regression baseline correction (see supplementary materials for details). Similarly, in contrast to our previous study, meditators in the current dataset did not show more frontal P3 topographies when activity was averaged from 416 to 512ms (*p* = 0.770), and meditators did not show more equal P3 amplitudes between Go and Nogo trials when global response strength was averaged from 336 to 449ms (*p* = 0.392). This was the case even when data were high pass filtered at 0.1Hz in closer replication of our previous study (both *p* > 0.05).

To further interrogate these potential inconsistencies between the studies by leveraging the increased power from a larger sample size, we performed an analysis of the easy Go/Nogo task data by combining data from both studies (after pre-processing all EEG data using the improved methods reported in the current study). The TANOVA analysis (of the distribution of activity) showed a main effect of group in the N2 window (from 191 to 244ms; global duration control = 49ms, *p*-value averaged across period of significance = 0.004, *np^2^* = 0.022), where meditators showed more negative voltages in frontal regions, providing further support for a stronger fronto-central N2 in meditators shown by our primary analysis of the age-matched groups in the data from the current study. A main TANOVA effect of group was also present in the P3 window (416 to 512 ms), with meditators showing more frontal P3 activity, similar to the results of our previous study.

However, the effect was small, and only just passed the statistical threshold (*p* = 0.047, *np^2^* = 0.002). We also note that the effect did not independently replicate with statistical significance in our new study data, rather it was only when the two studies were combined that the effect was significant.

Finally, the GFP test of global neural response strength showed a significant interaction between group and Go/Nogo trial type in the P3 window (386 to 429ms; global duration control = 43ms, *p*-value averaged across the period of significance = 0.007, *np^2^* = 0.062). This effect was driven by meditators showing larger Nogo P3 GFP values than non-meditators (as well as larger Nogo P3 GFP values than their Go trial P3 GFP values, while non-meditators showed no differences between Go and Nogo trials). This effect occurred within a similar window to the group by Go/Nogo trial type interaction reported in our original study. However, in the previous study, non-meditators exhibited larger Go trial P3 GFP compared to Nogo trials, whereas meditators showed no difference, which is almost the opposite of the pattern observed here.

To further assess this discrepancy, we repeated our analysis after 1) filtering data with a 0.25 Hz high-pass filter, and 2) applying a subtraction baseline correction approach. The result was replicated when we high pass filtered our data at 0.25 Hz, indicating the 0.5 Hz high-pass filter setting in the new study was not responsible for the altered effect compared to the original study (*p*-value averaged across the period of significance = 0.003, *np^2^* = 0.066). However, when we used the subtraction baseline correction method, the result of this exploratory analysis was no longer significant (*p* = 0.301, *np^2^* = 0.010). Furthermore, the subtraction baseline correction data showed a similar pattern to the results reported in the original study, with meditators showing less difference between the Go and Nogo trials than non-meditators. This suggests that our original result was likely driven by the confounding influence of a subtraction baseline correction method, which can project differences from the baseline correction period into the active period (Alday, 2019; Bailey et al., 2023c; Bailey et al., 2023e).

## Discussion

In this study, we aimed to determine whether neural activity related to conflict monitoring, expectation, and attention processes showed different characteristics in experienced mindfulness meditators, relative to non-meditating controls. Our results indicated that the fronto-midline N2 response, an ERP related to conflict monitoring and stimulus expectations, was larger in meditators in response to Nogo trials in a hard version of the Go/Nogo task. Evidence from age-matched comparisons and single trial analyses also indicated that the fronto-midline N2 was larger in meditators across both Go and Nogo conditions, and in the easy and hard version of the task. Our single trial analysis further indicated that more pronounced fronto-midline N2 responses were related to correct non-responses to Nogo trials, possibly reflecting enhanced attention to stimuli. This suggests that the enhanced fronto-midline N2 observed in meditators may represent attention-driven adaptive neural activity that enables better task performance.

Our results further revealed that meditators exhibited a more posteriorly distributed P3 response in hard Nogo conditions, and a more frontally distributed Nogo P3 response in easy conditions. Conversely, non-meditators exhibited a P3 response in hard Nogo conditions that was more frontally distributed compared to their easy Nogo P3. Our single trial analyses suggests that a more frontal P3 distribution is likely related to both stronger top-down expectations for Go trials (in the hard task) and to the allocation of top-down attention to sensory information (in the easy task). More frontal Nogo P3 responses were also associated with a higher proportion of correct responses, with a stronger effect in the easy task compared to the hard task. This suggests that meditators’ more frontal Nogo P3 activity in the easy task may reflect an attentional mechanism supporting higher task accuracy. Finally, while there was no group difference in overall task accuracy, a comparison restricted to age-matched groups indicated that meditators performed the easy (but not hard) task more accurately, in replication of our previous study (Bailey et al., 2019a).

Overall, the results suggest that meditation experience is associated with several differences in brain activity during the Go/Nogo task. While the altered ERPs in meditators are associated with conflict monitoring and stimulus expectation, the meditation-related effects are more likely driven by attentional differences rather than processes specific to conflict monitoring or stimulus expectation.

### Meditators Show Stronger Fronto-Midline N2 Activity

To enable us to interpret the finding that meditators exhibited stronger fronto-midline N2 activity, an understanding of the functional relevance of potential differences in the N2 is critical. One interpretation suggests that the N2 indexes the processing of mismatches (prediction errors) between higher-order or “global rule” aspects of the brain’s predictive model and sensory information (Schröger et al., 2015). In this context, prediction errors are greater for Nogo trials than Go trials (due to a response tendency), and greater for hard Nogo trials than easy Nogo trials (due to higher expectations for Go trials). However, interpretation of the functional relevance of the N2 must account for the smaller N2 amplitude as the number of prior Go trials accumulated in the hard version of the task (for both Go and Nogo trials). One potential explanation is that the repetition of Go trials in the hard task progressively weakened stimulus expectations based on participants’ perception of global stimulus frequencies, resulting in a N2 that was engaged slightly less strongly after higher numbers of Go trials in the hard task. This effect appeared to plateau after approximately six trials, possibly due to the influence of local frequencies on participant expectations overwhelming global expectations. In contrast, in the easy task, the global rule required 50% of trials to be Nogo trials; hence, participants may have increasingly expected a Nogo trial as the number of prior Go trials approached the point where additional Go trials would be improbable based on this global rule, thus explaining the accompanying reduction in Nogo N2 amplitudes with increasing numbers of preceding Go trials. This suggests that the fronto-midline N2 also reflects the processing of sensory information based on stimulus expectations derived from the interaction between local expectations (influenced by trials immediately preceding the current trial) and global expectations (based on overall stimuli probabilities) in a way that is modulated by attention (Cavanagh et al., 2010; Lauffs et al., 2020; Wacongne et al., 2011).

Given this pattern of results, our findings have several potential implications. First, our exploratory analyses suggest the fronto-midline N2 likely reflects cognitive processes beyond conflict monitoring, in alignment with previous research (Smith et al., 2010). In the harder task, and in Nogo trials, our results did show that the fronto-midline N2 was larger in amplitude, supporting its role in processing conflict monitoring, consistent with previous research (Donkers & Van Boxtel, 2004; Falkenstein, 2006; Smith et al., 2010).

However, our findings suggests that the fronto-central N2 can be modulated by attention effects as well as conflict monitoring effects. In this context, the stronger fronto-midline N2 amplitude in meditators may indicate attention-related increased processing of sensory information, an effect that may interact with task demands to be larger when conflict monitoring requirements increase, such that the differences in meditators were most detectable in hard Nogo trials. We conjecture that this effect may be related to meditators’ attentional training (Crane et al., 2017; Van Dam et al., 2018). In contrast, non-meditators showed lower fronto-midline N2 amplitudes in the hard Nogo trials, perhaps reflecting more habitual responses and less attention to sensory information.

These results aligns with previous research identifying structural changes in the ACC in meditators, with this brain region thought to generate the fronto-midline N2 (Allen et al., 2012; Falcone & Jerram, 2018; Ganesan et al., 2022; René J Huster et al., 2013; Tang et al., 2010; Tomasino et al., 2016). They also highlight the likely necessity for a nuanced understanding of how attention and executive control functions are modified by mindfulness meditation. This nuanced understanding requires further interrogation in future research, and we note that our current interpretation of the meditation-related differences in the N2 response remains speculative. However, it does align well with the present findings. An alternative explanation might be that meditators formed stronger expectations about the upcoming stimuli, resulting in a larger conflict between expectation and the stimuli, thereby eliciting a larger amplitude fronto-midline N2 to resolve the conflict. However, while this explanation fits with the enhanced fronto-midline Nogo N2 in meditators, it does not align with the relationship between enhanced fronto-midline N2 activity and correct responses, as increased expectations for Go trials could be assumed to be associated with more incorrect responses to Nogo trials.

#### Meditators and Non-Meditators Showed Different Patterns of P3 Activity Across the Easy and Hard Tasks

Our analysis indicated that meditators showed a more frontally distributed P3 in the easy task compared to the hard task, with the effects of task difficulty in meditators being strongest across frontal electrodes. In contrast, non-meditators showed a more frontally distributed P3 in the hard task compared to the easy task, with task difficulty effects in non-meditators appearing most strongly in electrodes over parietal regions. To understand this pattern of results, we analysed how the P3 centroid was modulated by different task demands, including task difficulty, response tendencies and response inhibition requirements, as well as stimulus expectations based on both task difficulty and local stimulus frequencies. Our exploratory analysis of the P3 suggests that its degree of frontal distribution was related to task difficulty, the number of prior Go trials, and whether a trial was a Go or Nogo trial. As such, the distribution of the P3 seems to be affected by both local and global stimuli probabilities, perhaps reflecting the processing of prediction errors related to expectations about upcoming stimuli, as well as cognitive control, response inhibition and likely attention processes, in alignment with functional interpretations of the P3 provided by previous research (Clayson & Larson, 2011; Datta et al., 2007; Kluger et al., 2020; Smith et al., 2008; Wickens et al., 1983).

One potential explanation of the P3 topography effects that accounts for all aspects of our pattern of results is that the degree of frontal distribution of the P3 might be modulated by both the strength of top-down expectations (as per the response tendency associated with an expectation for Go trials), but also by the allocation of top-down attention to sensory information to detect “informative events”, which causes prediction errors to be passed to higher levels of the cortical hierarchy (Kluger et al., 2020). The more frontal P3 topography might reflect the allocation of additional resources to process these informative events to resolve uncertainty, an effect that can cause increased stimulus processing even when the stimulus does not reflect a prediction error (Kluger et al., 2020). Within this interpretation, an increasing number of Go trial repeats might increase the participant’s uncertainty about the stimulus type in upcoming trials, particularly in the easy task where lower global frequencies of Go trials were expected (a maximum of four consecutive Go trials were presented in the easy task, which may have enabled participant responses to be informed by order effects). The processing of these higher order sequence effects has been shown to be performed in

higher regions of the neural hierarchy (Wacongne et al., 2011). Heightened uncertainty in specific task conditions may promote increased attention to the stimulus, enabling more precise processing of this stimulus information, greater transmission of prediction errors to higher cortical areas, and more prediction model updating. As such, increased attention to resolve increased uncertainty might explain the increased frontal distribution of the P3 with more preceding Go trials, irrespective of whether a Go or Nogo trial elicited the P3. This effect appears to be independent of frequency-based expectation violations produced by a Nogo trial in the hard task, or a response-tendency-based expectation violations produced by a Nogo trial in the easy task. This explanation also accounts for the association between more frontally distributed P3 responses and correct non-responses to Nogo stimuli, with trials containing more frontal P3 activity reflecting more attention to the task, enabling an attention-based performance increase.

This pattern may explain why the frontal shift in the P3 with increasing numbers of prior Go trials was more pronounced in the easy task than the hard task. It suggests that prediction errors were processed higher in the cortical hierarchy when the local stimulus probabilities preceding the current trial were less aligned with global stimulus probabilities. This conflict between local and global stimulus frequencies may be associated with increased uncertainty in the brain’s predictive model, provoking engagement of attentional resources to process informative events in an attempt to resolve uncertainty. This increase in attention engagement associated with elevated processing of informative events for the brain’s predictive model may take place irrespective of whether the events are aligned with predictions (Kluger et al., 2020).

The potential gains in task performance from awareness of higher order sequence effects were reduced in the hard task, potentially diminishing the functional relevance of a more frontal P3 response in this condition. Based on this interpretation, we propose that meditators’ stronger engagement of frontal regions in the easy condition may represent an attentional mechanism that supports the processing of informative events at higher levels of the neural hierarchy, possibly facilitating their better task performance in the easy task. Conversely, the meditators’ less frontally distributed P3 response in the hard task may reflect more weakly held predictions about upcoming stimuli, leading to reduced engagement of frontal brain regions in processing prediction errors, and perhaps the engagement of stimulus driven ‘bottom-up’ attention underpinned by posterior regions to process stimuli in the hard task (Zhang et al., 2022).

In contrast, the task difficulty-based difference in P3 activity was strongest in posterior regions for the non-meditators, suggesting that they did not engage top-down attention mechanisms to process stimuli more precisely in the easy version of the task. Furthermore, non-meditators showed a P3 that was more frontally distributed in the hard version of the task. This might reflect an increased generation of top-down predictions in the hard task, and an associated increase in prediction error processing in higher cortical regions when higher Go stimulus frequencies generated stronger expectations for upcoming Go trials. Since the more frontal P3 responses in the hard task in non-meditators were likely driven by an increased mismatch between predictions and the Nogo stimuli rather than an increase in attention, these more frontal P3 topographies in the hard task were present without the performance gains based on the allocation of attention to stimuli shown by the meditators in the easy task. In support of these interpretations, engagement of frontal regions during the P3 was associated with a higher proportion of correct responses across both groups, and this effect was stronger for the easy task than the hard task.

These results are conceptually aligned with previous research showing that P3 activity mediates the relationship between high trait mindfulness and high executive function performance (Lin et al., 2019). However, given our current results did not directly replicate our previous research, they should be interpreted with caution. It is also worth noting that a null result for both the N2 and P3 in long term meditators has been reported by previous research examining the Go/Nogo task (Andreu et al., 2019). In that study, the task was easier than the hard task reported in our study (with a higher Nogo frequency), and single electrode analyses were implemented, which are not sensitive to differences in the distribution of activity (which were the primary results in the current study). Nonetheless, additional research is required to replicate our results before we can determine whether our interpretations of the relationship between meditation and neural activities are correct.

### Limitations and Future Directions

As is typical in EEG research, we made specific choices about how to pre-process our EEG data prior to analysis. We based these choices on current best practice, using objective methods that minimised potential confounds and maximised signal to noise ratios (Bailey et al., 2023b; Bailey et al., 2023c). We also tested whether our results were robust to a number of permutations in the pre-processing parameters as is recommended best practice (Bailey et al., 2023b; Bailey et al., 2023c). As such, we consider that our results are robust to specific analytical choices. However, we note that EEG pre-processing methods are continually developing (Bailey, Hill, Godfrey, Perera, Rogasch, et al., 2024), and future research may provide a more optimal approach. Additionally, we note that our study was cross-sectional in design, so does not enable inferences about causation.

Interestingly, and in contrast to our expectations that the hard version of the task would be more sensitive to group differences in task accuracy, the easy version of the task showed differences between the two groups in task accuracy when analyses were restricted to age-matched groups. This result replicates our previous finding that meditators performed better in the easy version of the Go/Nogo task, as well as other research showing higher Go/Nogo task performance in meditators (Andreu et al., 2019; Sahdra et al., 2011; Zanesco et al., 2013). It may be that the easy version of the task was monotonous, and so reflected a sustained attention challenge, with the meditation group able to maintain task focus and show better performance, while the non-meditator group suffered from inattention and performed slightly worse. This finding may be highly relevant for future research attempting to characterise the effects of meditation on cognition. Given this possibility, incorporation of a sustained attention task in future research could be useful to better disentangle any such effects. Current research has shown mixed results for the effects of meditation on cognition, with overall small effect sizes reported in meta-analyses (Sumantry & Stewart, 2021; Yakobi et al., 2021). It may be that any cognitive effects related to meditation are dependent on specific circumstances. We note here that while there is considerable research on the cognitive effects of meditation, the landscape of those effects across different task parameters is currently not fully characterised.

## Conclusion

Overall, our findings demonstrate that meditators perform more accurately in an easy Go/Nogo task (but not a hard Go/Nogo task), and that their improved performance is likely to be underpinned by attention-related changes in the distribution of neural activity, which perhaps reflect increased engagement of frontal brain regions to process sensory information with more precision. These results lend further support to the suggestion that meditation has neuroplastic effects on the brain, and that the effects reflect attention-related mechanisms, even when the neural activities being measured reflect the engagement of conflict monitoring and stimulus expectation-based processes.

## Acknowledgements

We gratefully acknowledge the Dhamma Aloka Vipassana meditation centre in Melbourne and the Melbourne Insight Meditation centre for their assistance with the recruitment of several meditators who took part in the study.

