## Supplementary Material for "Neural differences in conflict monitoring and stimulus expectancy processes in experienced meditators are likely driven by enhanced attention"

### Supplementary Materials

#### EEG pre-processing and analysis

The RELAX (Reduction of electroencephalographic artefacts) software was used to clean the EEG data (Bailey et al., 2023b; Bailey et al., 2023c). This pipeline firstly bandpass filters the data between 0.25Hz and 80Hz, with a notch filter from 47 to 53Hz, using fourth order Butterworth filters. The pipeline then uses the PREP toolbox to reject bad electrodes (Bigdely-Shamlo et al., 2015), followed by a secondary rejection of electrodes that show outlying data for more than 5% of the recording using established outlier identification approaches, with the limit that no more than 20% of electrodes could be rejected (Bailey et al., 2023b). Outlying periods remaining in the data were then rejected based on the same outlier identification approaches as the channel rejection step, then PREP's robust average re-referencing was applied to the data (Bigdely-Shamlo et al., 2015). Independent component analysis was then computed on the data using the PICARD algorithm (Frank et al., 2022). Artifactual components in the data were identified by the machine learning algorithm, ICLabel, with the criteria that if ICLabel identified a component as "most likely" to be artifact, it was selected for reduction (Pion-Tonachini et al., 2019). Artifacts from all categories were identified (eye movements/blinks, muscle activity, heart-beat, line noise, and non-specific other artifacts). These artifacts were then reduced using wavelet enhanced ICA (Castellanos & Makarov, 2006) before data were reconstructed back into the scalp space. Missing electrodes were interpolated back into the data using spherical spline interpolation (Perrin et al., 1989) prior to statistical analysis using the randomised graphical user interface (RAGU) (Koenig et al., 2011). A second set of the data were kept without interpolation for generalised eigenvector decomposition (which does not require interpolation of missing electrodes) (Cohen, 2022).

After cleaning, visual inspection of the data and the outcome metrics provided by RELAX suggested that a minority of files still showed either low frequency (<0.5Hz) drift or had muscle activity remaining. To address this, we bandpass filtered the data from 0.5 to 30 Hz prior to analysis. While high-pass filter settings have been debated for ERP analyses, a 0.5Hz high-pass filter (or even higher) has been recommended in some cases as providing improved detection of experimental effects (Bailey et al., 2024; Delorme, 2023) and is particularly helpful when slow drift is present in the data. In particular, we deemed that decreasing the influence of drift remaining in the data by using this setting was likely to lead to more reliable results for our GED decompositions (which rely on clean data) (Cohen, 2022) and in our single trial linear mixed model analyses (which can be adversely affected by outliers and measurement errors which can be produced by artifacts) (Bartlett et al., 2009). We conducted our analyses with RAGU and the GED ANOVA using both the 0.25Hz and 0.5Hz filter settings, and note that our results remained unchanged regardless of the setting used (although the effect sizes were smaller when applying the 0.25Hz high pass filter settings, likely as a result of the increased measurement noise with the 0.25Hz high pass filter settings). We also note that testing the 0.25Hz settings did not alter our results with regards to replicating our previous study, demonstrating that the filter settings were not responsible for replication or non-replication of specific results.

To perform the regression baseline correction, RELAX firstly averages the amplitude of voltage within the baseline period for each trial separately (Bailey et al., 2023c). The regression baseline correction is then performed on each channel separately by using a regression model (which includes the mean baseline activity, the Go and Nogo conditions and the hard and easy tasks as factors, along with a constant) to compute the standardized regression coefficients for the relationship between the mean baseline amplitude and the data at each timepoint across epochs. The mean baseline amplitude within each epoch is then multiplied by these standardized regression coefficients to obtain a model of how the baseline period for each epoch has influenced each time point within each epoch. The values from this baseline confound model are then subtracted from the epoched data to remove the baseline period's influence on the active data for each epoch, without transposing effects within the baseline period onto the active periods.

For our primary analyses, we used RAGU to make comparisons between groups and conditions. RAGU performs a GFP test to assess differences in global neural response strength by obtaining the GFP from 5000 randomisations of the data, shuffling individual maps between conditions/groups (which destroys any between condition/group effect in the real data) then ranking the size of the real GFP difference between conditions/groups against the shuffled data, with real data exceeding 95% of the shuffles deemed significant at $p < 0.05$ (Koenig et al., 2011). The TANOVA test calculates the average scalp-based map of activity for two separate conditions or groups, then subtracts the average of one condition/group from the average of the other condition/group (Koenig et al., 2011). This creates a global dissimilarity map. The GFP of this global dissimilarity map is then computed. Statistical comparisons are performed by obtaining the global dissimilarity map from 5000 randomisations of the data, shuffling individual maps between conditions/groups (which destroys any between condition/group effect in the real data), and ranking the size of the real GFP of the global dissimilarity map against the shuffled data, with real data exceeding 95% of the shuffles deemed significant at $p < 0.05$ (Koenig et al., 2011). Within the TANOVA test, GFP can be normalised to be equivalent to 1 for all participants prior to the TANOVA using a recommended L2 normalisation (Koenig et al., 2011). This enables the TANOVA comparisons to be independent of differences in global neural response strength, thus testing for a significant difference in the distribution of neural activity (reflecting differences in brain regions engaged) without the influence of differences in the strength of those activations. We used this L2 normalisation in our TANOVA analyses. Finally, to ensure that the groups and conditions tested showed within group consistency, we implemented the topographical consistency test, which uses randomisation statistics to compare the mean GFP to 0 (as per a single sample t-test design), ensuring that the assumption of within group topographical consistency is met before between group analyses progress. Figure S1 shows that the assumption of topographical consistency was met for all groups, conditions, and almost all timepoints in the epoch (with the exception of a brief period that did not overlap with any of our significant effects of interest in our primary analyses).

**Generalised Eigenvector Decomposition and Single Trial Analyses**

To understand the functional significance of the differences we detected in our primary analyses, we performed generalised eigenvector decompositions to spatially filter the data separately for each individual and extract source space phasic ERP signals that reflected the topographical differences shown by RAGU to differ between the groups (Cohen, 2022). We then performed single trial analyses implemented with linear mixed model analyses and generalised linear mixed model analyses to interpret the functional relevance of these GED decomposed neural activities that differed between the groups where possible. However, while the N2 effect provided a clear scalp map target for GED extraction, a P3 difference did not. As such, single electrode ERPs average during the significant effect involving group were used in the linear mixed models instead.

To achieve these GED decompositions, first, we obtained an average ERP across all electrodes for each Go/Nogo and easy/hard task condition separately using timepoints from 150ms to 600ms (reflecting the N2 and P3 activity). We then used this ERP map to construct the signal covariance matrix. Following this, we constructed the reference covariance matrix from all single trial data (without averaging the reference data, so the reference data contained the phase locked data and also the non-phase locked activity). Both signal and reference signals were computed after excluding outlying single trial epochs with z-scored Euclidean distances in the covariance matrices of more than 2.31 (corresponding to $p < 0.01$). The reference covariance matrix was then regularised by shrinking by 1% of the amplitude from all datapoints to improve matrix separability (Cohen, 2022), and the signal covariance matrix was normalised to the amplitude of the reference covariance matrix. The GED was then used to obtain a list of spatial weighting components and eigenvalues reflecting the maximal separation between the reference and signal covariance matrices (the maximal phasic ERP activity).

To resolve sign uncertainty (where the GED produces time courses without polarity necessarily matching the scalp space) we correlated the GED component time course with the ERP from electrode FCz (where the N2 was often largest), and inverted the GED polarity if the correlation was negative (Cohen, 2022). Then, to extract the fronto-midline N2 component (which our primary analyses with RAGU indicated to differ between groups), we constructed a scalp electrode map with a gaussian distribution of positive/negative voltages based on a maximum at FCz and diminishing from positive to negative voltages with increasing distance from FCz (Zuure et al., 2020). We extracted the largest GED component that showed a squared Pearson's correlation ($R^2$) > 0.45 between the GED component and the fronto-midline activity template (reflecting high shared spatial variance) from each individual for each condition (Zuure et al., 2020). We examined a GED extracted fronto-midline N2 component averaged during the 150 to 250ms N2 window. This window was established prior to further analysis by both considering the 198 to 241ms period of significant 3-way interaction in the RAGU analysis, and by inspecting the GED N2 deflection in the grand averaged data across all groups and conditions, which showed a distinct peak onset at 150ms and offset at 250ms (see Figure 4 in the main manuscript). Averaged values from within this 150ms to 250ms period were submitted to the linear mixed model analyses and generalised linear mixed model analyses.

To understand the potential functional relevance of the differences in N2 and P3 activity in the meditators, we conducted a number of linear mixed model and generalised linear mixed model analyses based on single trials from the correct responses to easy and hard Nogo trials. Linear mixed model (LMM) analyses were computed using JASP 0.17.2.1. (using the Satterthwaite test method type III). The linear mixed model analyses included as dependent variables either: 1) the mean centred and z-score corrected values from the fronto-midline GED component time course averaged between 150 to 250ms after stimulus presentation, or 2) the centroid of the topographical distribution of the P3, obtained by multiplying the x coordinates of all midline electrodes by the averaged voltage at each electrode within the 339 to 405ms period after stimulus presentation (the period that showed significant effects in our TANOVA analyses), weighted by the absolute averaged voltage value from each electrode during the same period. This indicated the extent of the frontally distribution of the P3, with more positive values (up to a maximum of 85 for electrode FPz) reflecting more frontal distributions, and more negative values (down to a minimum of -85 for Oz) reflecting more posterior distributions. For these LMM analyses, participant was included as the random effects variable. Group, Nogo difficulty (easy / hard Nogo trials), and the number of prior Go trials were included as fixed effects variables. For analyses that included Nogo trials from the easy task and number of prior Go trials as a factor, we restricted the number of prior Go trials to a maximum of 4 so that the easy and hard versions of the task were matched. Similarly, for analyses that included Go trials from the easy task, we restricted our analyses to a maximum of 3 prior Go trials (as the easy task presented a maximum of four consecutive Go trials, such that analysis of the Go P3 could include a maximum of 3 prior Go trials). To understand the effects of higher numbers of repeated prior Go trials, we also performed analyses that only included the hard task and restricted the number of prior Go trials to a maximum of 9. The task design included all numbers of preceding Go trials from 0 to 9, but due to the randomisation of trial order, had intermittent gaps in the number of preceding Go trials between 10 and 18, so restricting to a maximum of 9 prior Go trials ensured all participants provided representation at each included level in the analysis. When singular model fits occurred, we reduced the complexity of the analysis by a single factor at a time then re-computed the analysis until the model did not report a singular fit.

In addition to the LMM focused on examining the effects of group, task difficulty, and the number of prior Go trials on the N2 and P3, we used generalised linear mixed model (GLMM) analyses to examine how the N2 or

P3 activity affected the likelihood of accurate responses, either across both tasks or in either the hard Nogo trials or the easy Nogo trials separately. We also used a GLMM analysis to test whether the relationship between the ERP and response accuracy was modified by number of prior Go trials and task difficulty. For all these analyses, participant was included as the random effects variable. Group, task difficulty (easy / hard Nogo trials), and the number of prior Go trials were included as fixed effects variables. To avoid singular model fitting from overly complex analysis design, we reduced the fixed effects variables included in the model where singular fits occurred and tested different conditions in separate analyses to assess potential interactions where singular fits occurred.

#### Supplementary Materials Results

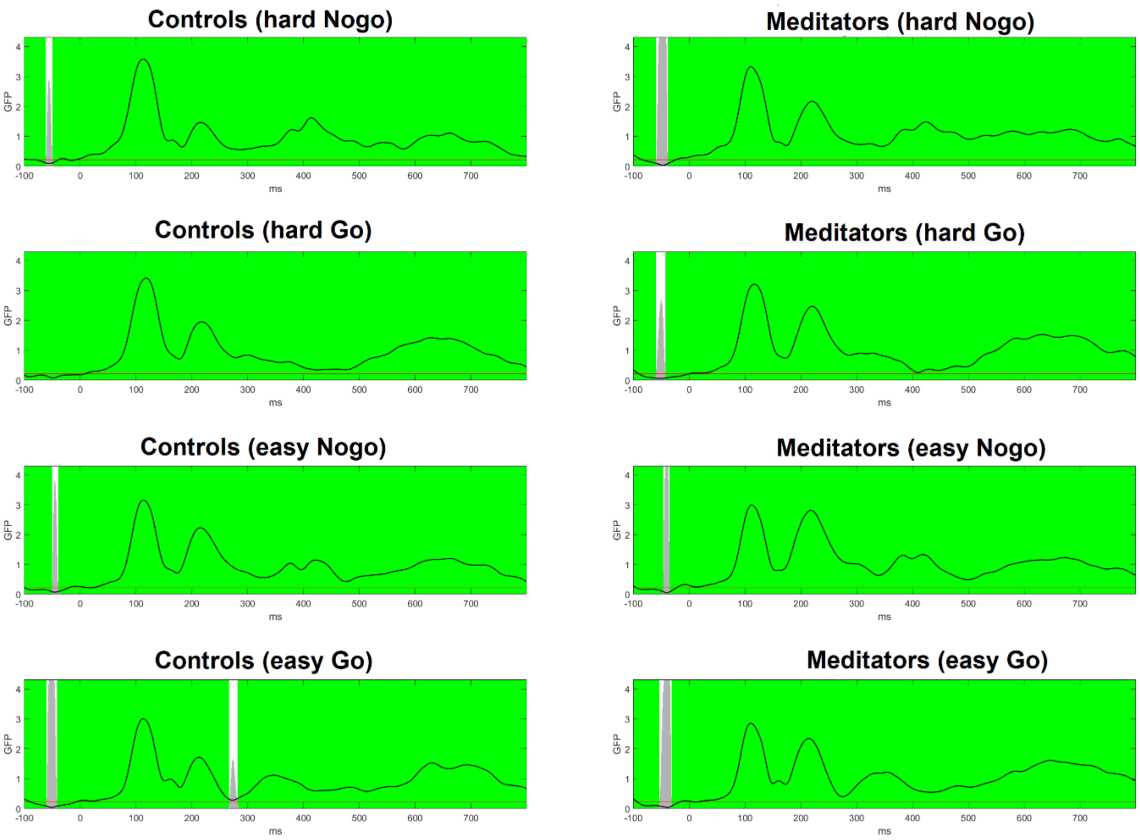


**Figure S1.** Topographical consistency tests demonstrated consistent neural activity distribution patterns in both groups and all conditions, with the exception of a brief period just prior to 300ms in the easy Go trial condition for non-meditators (controls). Note that this period did not overlap with significant periods for effects involving group detected by RAGU. The black line reflects the global field potential computed from the average voltage at each electrode across each participant within the condition (this GFP value would equal zero if there were no consistent pattern in the distribution of neural activity). Green periods reflect periods showing consistent activity, grey bars reflect the *p*-value (which is below the *p*-value of 0.05 for most of the epoch in all conditions), the red line at the bottom reflects the $p < 0.05$ threshold, and white periods reflect periods showing inconsistent activity.

#### GED and Linear Mixed Modelling Analyses

*<u>LMM of N2 GED analysis of correct responses to Nogo trials:</u>*

Linear mixed-effects modelling of trial-level N2 amplitude (as estimated using generalised eigenvector decomposition) was performed to investigate how N2 responses on Nogo trials differed across non-meditator controls and experienced meditators as a function of the number of preceding Go trials in both the easy and hard Go/Nogo tasks. The number of prior Go trials was constrained to a maximum of 4 in the easy task and 9 in the hard task; this factor was modelled using $1^{st}$- and $2^{nd}$-degree polynomial terms to account for potential curvilinear effects. Random slopes for the number of prior Go trials and task difficulty were included on random intercepts for subject identity.

Model comparison confirmed that the inclusion of $2^{nd}$-degree polynomial terms was justified by a significant improvement in model fit compared to a reduced model that included only $1^{st}$-degree (i.e., linear) terms

($\chi^2(8) = 44.37$, $p < 0.001$). Type-II analysis-of-variance revealed a significant main effect of group, whereby the N2 component was significantly more negative in meditators than in control subjects ($F(1,62) = 6.59$, $p = 0.013$). There was also a significant main effect of prior Go trials ($F(2, 57.3) = 4.14$, $p = 0.021$), and a significant two-way interaction between the number of prior Go trials and task difficulty ($F(2, 8837.5) = 28.30$, $p < 0.001$). While N2 amplitude in the easy Nogo task remained reasonably stable as the number of prior Go trials increased from 0 to 4 (nadiring at 2 Go trials), it became progressively less negative in the hard Nogo task, remaining relatively stable thereafter (see Figure S2). There was no significant interaction between number of prior Go trials and group ($F(2,57.9) = 1.08$, $p = 0.347$) or between group and task difficulty ($F(1,62) = 3.95$, $p = 0.051$); the three-way interaction between all factors was also non-significant ($F(2,8839) = 1.39$, $p = 0.249$).

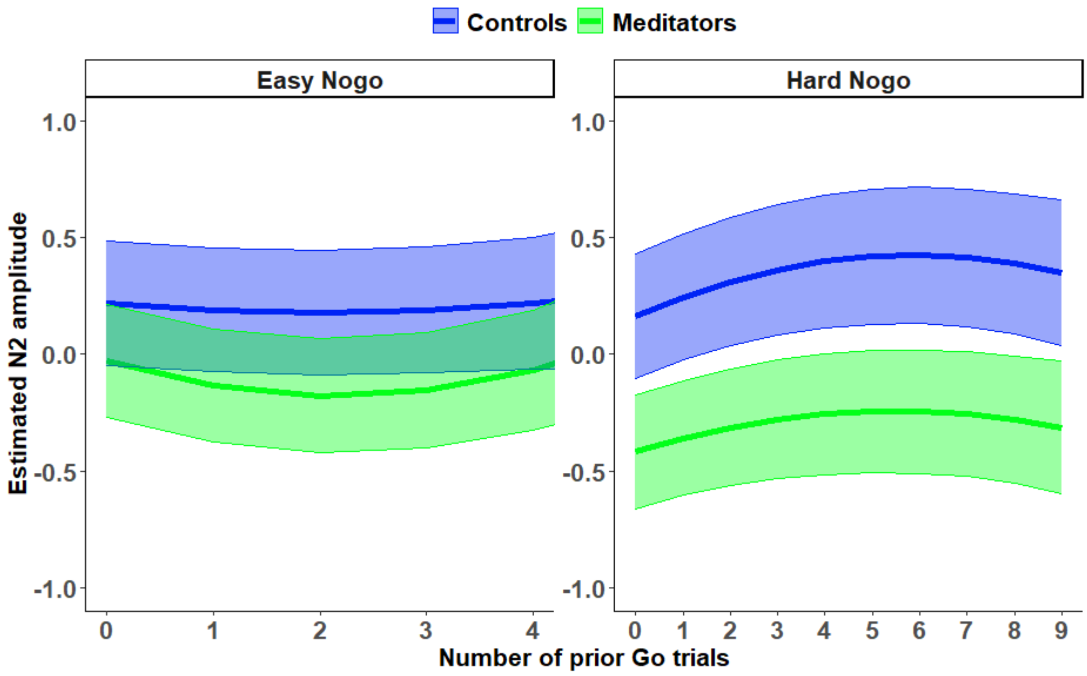


**Figure S2.** Estimated marginal mean GED N2 component amplitudes for Nogo trials as a function of the number of prior Go trials and task difficulty (easy vs. hard). Error shades reflect 95% confidence intervals.

*LMM of N2 GED analysis of correct responses to Go trials:*

To better understand the functional relevance of the N2 component, we refit the LMM reported above to N2 components following Go trials. Again, this model revealed a significant main effect of group, whereby GED N2 component amplitude was more negative in meditators versus controls ($F(1,62) = 6.14$, $p = 0.016$). There was no significant main effect of either the number of prior Go trials ($F(2,60.6) = 2.23$, $p = 0.116$) or task difficulty ($F(1,62) = 0.15$, $p = 0.700$). While the between-group difference in N2 amplitude appeared more pronounced in the easy task than the hard task (see Figure S3), the two-way interaction was non-significant ($F(1,62) = 0.88$, $p = 0.352$). There was however a significant interaction between task difficulty and number of preceding Go trials ($F(2,20424) = 12.49$, $p < 0.001$). While N2 amplitude tended to become more negative as the number of prior Go trials increased in the easy task, it became slightly less negative as the number of prior Go trials increased in the hard task. This effect is reminiscent of the two-way interaction observed for Nogo trials. As with the Nogo trial analysis, there was no significant interaction between number of prior Go trials and group ($F(2,60.7) = 0.69$, $p = 0.508$), or the three-way interaction between all factors ($F(2,20427) = 2.18$, $p = 0.113$).

Overall, these results demonstrate that the N2 varies in response to interactions between global frequencies (across both tasks) and local frequencies (with similar patterns across both Go and Nogo trials, suggesting attention engagement effects), as well as conflict monitoring / response inhibition demands.

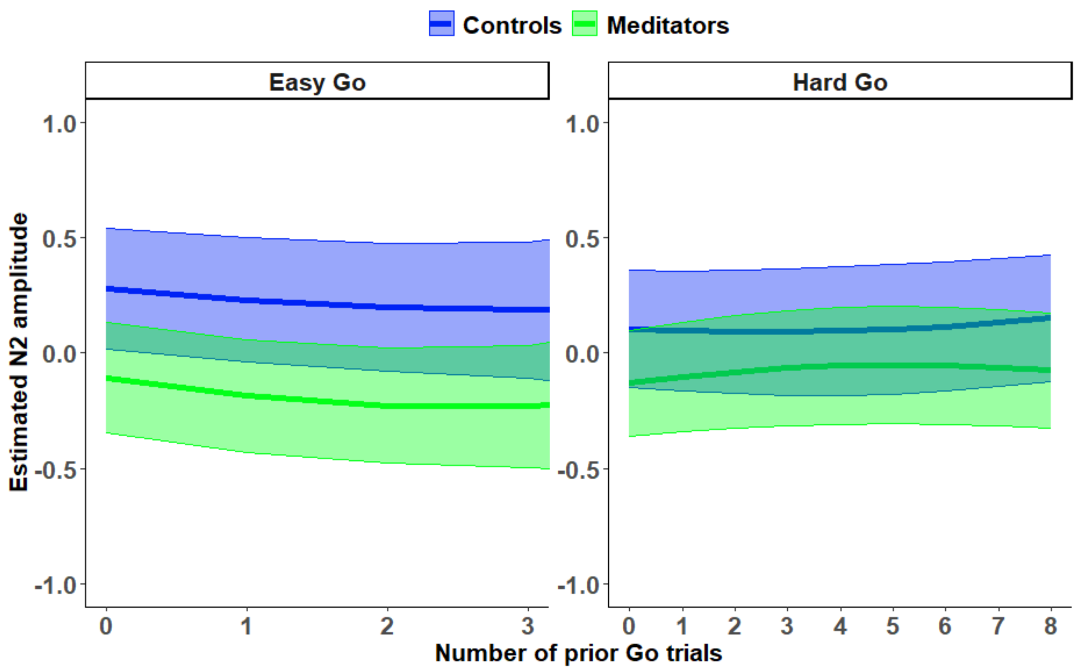


**Figure S3.** Estimated marginal mean GED N2 component amplitudes for Go trials as a function of the number of prior Go trials and task difficulty (easy vs. hard). Error shades reflect 95% confidence intervals.

*<u>GLMM of N2 GED analysis of accuracy of responses to Nogo trials:</u>*

When both the easy and difficult tasks Nogo GED N2 amplitudes were included in a GLMM analysis, but task difficulty was not included as a factor, there was no relationship between N2 component amplitude and correct responses (*ChiSq* = 2.323, *p* = 0.127). Unfortunately, a singular fit occurred for the interaction between task and Nogo GED N2 amplitude, so the tasks were tested separately. The results for the analysis of the hard Nogo trials as the dependent variable indicated a significant main effect of the Nogo GED N2 component amplitude on the likelihood of correct response (*ChiSq* = 8.950, *p* = 0.003), where larger (more negative) amplitudes were associated with more correct responses (see Figure S4 and Table S1). This result was significant even when the number of prior repeated Go trials was restricted to a maximum of 4 (to match the prepotent response demands present in the easy Nogo trials) (*ChiSq* = 9.589, *p* = 0.002). In contrast, within the easy task there was no relationship between Nogo GED N2 amplitude and the likelihood of correct responses (*ChiSq* = 0.312, *p* = 0.577). When testing the relationship to correct responses in the hard task only, there was no significant interaction between Group and Nogo GED N2 component amplitude (*ChiSq* = 1.030, *p* = 0.310). Nor was this interaction significant in the easy task (*ChiSq* = 0.137, *p* = 0.711).

As expected, the number of prior Go trials did decrease the probability of a correct response as expected (*ChiSq* = 9.399, *p* = 0.002). However, there was also no interaction between the Nogo N2 component amplitude and the number of prior Go trials when hard Nogo correct/incorrect responses were the dependent variable (*ChiSq* = 0.138, *p* = 0.710). Nor was there an interaction between the Nogo N2 component amplitude and the number of prior Go trials when easy Nogo correct/incorrect responses were the dependent variable (*ChiSq* = 0.759, *p* = 0.384), and unexpectedly, an increase in the number of prior repeated Go trials did not decrease the probability of a correct response (*ChiSq* = 0.414, *p* = 0.520). This pattern of results suggests the relationship between the N2 amplitude and correct responses was driven by global Go and Nogo frequencies (where the hard task presented 25% Nogo trials overall) rather than by local frequencies (where an increasing number of preceding Go trials did not alter the relationship between Nogo N2 amplitudes and correct non-responses).

**Table S1.** Estimated marginal means for the GLMM including correct/incorrect response to the hard Nogo trials as the dependent variable, and Nogo GED N2 amplitude and group as fixed effect variables, with participant as the random effects variable. Note that results are on the response scale.

**Estimated Marginal Means**

| | | | | 95% CI | |
|---|---|---|---|---|---|
| **GED N2** | **Task** | **Estimate** | **SE** | **Lower** | **Upper** |
| -1.000 | Easy | 0.961 | 0.007 | 0.946 | 0.972 |
| 0 | Easy | 0.963 | 0.005 | 0.952 | 0.972 |
| 1.000 | Easy | 0.965 | 0.006 | 0.951 | 0.976 |
| -1.000 | Hard | 0.784 | 0.025 | 0.731 | 0.829 |
| 0 | Hard | 0.745 | 0.025 | 0.693 | 0.790 |
| 1.000 | Hard | 0.701 | 0.031 | 0.636 | 0.759 |

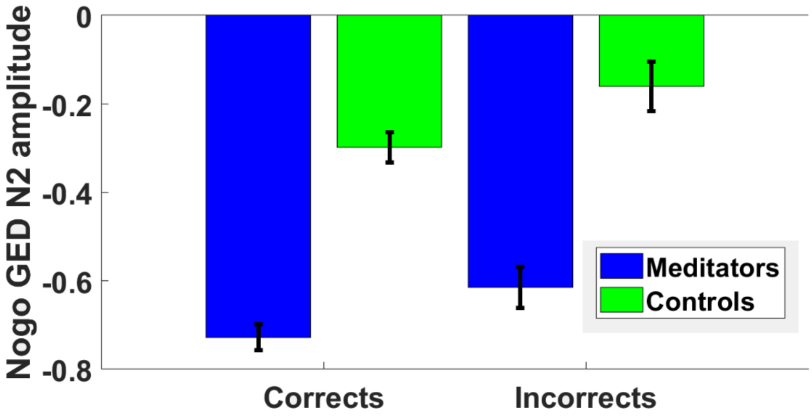


**Figure S4.** Mean Nogo GED N2 for correct and incorrect responses for the meditation and control group separately for the hard Nogo trials. Error bars reflect 95% confidence intervals.

*LMM analyses of the P3 centroid within the P3 window in correct Nogo trials:*

In addition to our single trial exploration of the N2, we also conducted a number of LMM analyses in an attempt to understand the functional relevance of the interaction between group and task difficulty in the topographical distribution of neural responses during the P3 window (339 to 405ms). The interaction involving group seemed to be driven by topographical differences between the groups in response to the different task difficulties. Unlike the N2, the GED approach did not extract topographies with maximums at these electrodes consistently across participants, so single electrode analyses were used, examining the P3 centroid to assess differences in the frontal distribution of the P3.

Our analysis of the Nogo P3 centroid showed an interaction between group and task difficulty ($F$(1,59.85) = 7.588, $p < 0.008$), with the meditators showing a more frontally distributed Nogo P3 in the easy task, while controls showed a more frontally distributed Nogo P3 in the hard task (Table S2). The Nogo P3 also shifted further forward as a higher number of preceding Go trial repeats had been presented ($F$(1,50.82) = 21.939, $p$ < 0.001), and an interaction effect was present where the forward shift in the Nogo P3 was stronger for the easy task than the hard task ($F$(1,62.65) = 10.277, $p$ = 0.002, Table S3).

**Table S2.** Estimated marginal means for the effect of different task conditions and group on the Nogo P3 centroid.

**Estimated Marginal Means**

| | | | | | 95% CI | |
|---|---|---|---|---|---|---|
| **Condition** | **Prior Go Trials** | **Group** | **Estimate** | **SE** | **Lower** | **Upper** |
| Easy | -0.129 | Non-meditators | 7.902 | 3.098 | 1.830 | 13.975 |
| Hard | -0.129 | Non-meditators | 14.264 | 3.210 | 7.973 | 20.555 |
| Easy | 1.102 | Non-meditators | 14.256 | 2.486 | 9.383 | 19.129 |
| Hard | 1.102 | Non-meditators | 17.378 | 2.891 | 11.712 | 23.043 |
| Easy | 2.334 | Non-meditators | 20.610 | 2.704 | 15.310 | 25.910 |
| Hard | 2.334 | Non-meditators | 20.491 | 3.365 | 13.895 | 27.086 |
| Easy | -0.129 | Meditators | 16.671 | 2.809 | 11.167 | 22.176 |
| Hard | -0.129 | Meditators | 16.165 | 2.889 | 10.502 | 21.827 |
| Easy | 1.102 | Meditators | 21.935 | 2.254 | 17.517 | 26.354 |
| Hard | 1.102 | Meditators | 16.328 | 2.599 | 11.234 | 21.422 |
| Easy | 2.334 | Meditators | 27.199 | 2.439 | 22.419 | 31.979 |
| Hard | 2.334 | Meditators | 16.491 | 3.000 | 10.611 | 22.371 |

**Table S3.** Estimated marginal means for the effect of different task conditions and group on the Go P3 centroid.

**Estimated Marginal Means**

| | | | | 95% CI | |
|---|---|---|---|---|---|
| **How Many Prior Go trials** | **Task** | **Estimate** | **SE** | **Lower** | **Upper** |
| -0.065 | Easy | -14.584 | 2.395 | -19.279 | -9.890 |
| 1.225 | Easy | -7.120 | 2.204 | -11.439 | -2.801 |
| 2.515 | Easy | 0.345 | 2.528 | -4.609 | 5.299 |
| -0.065 | Hard | -5.289 | 2.085 | -9.375 | -1.203 |
| 1.225 | Hard | -2.529 | 1.946 | -6.343 | 1.286 |
| 2.515 | Hard | 0.232 | 1.977 | -3.643 | 4.106 |

*GLMM analyses of relationship between Nogo response accuracy and P3 activity at single electrodes:*

In addition to the LMM, we used GLMM analyses to examine how the P3 centroid affected the likelihood of accurate responses in either the hard Nogo trials or the easy Nogo trials, and whether the relationship between the P3 and response accuracy was modified by number of prior Go trials and task difficulty. For all these analyses, participant was included as the random effects variable. Group, task difficulty (easy / hard Nogo trials), and the number of prior Go trials were included as fixed effects variables. In order to avoid singular model fitting from overly complex analysis design, we restricted the fixed effects variables included in the model to two in any single analysis.

There was a significant main effect of P3 centroid, whereby more frontal P3 centroids were associated with correct responses (*ChiSq* = 4.875, *p* = 0.027 (Table S6). There was also an interaction between task difficulty the P3 centroid, where the increase in correct responses for more frontal P3 centroids was larger in the easy task than the hard task (*ChiSq* = 6.072, *p* = 0.014).

**Table S6.** Estimated marginal means for the interaction between Nogo P3 centroid and task difficulty relative to correct responses. Note that in the easy task a more frontal Nogo P3 was associated with a larger increase in the proportion of correctly responded Nogo trials than when the Nogo P3 centroid was more frontal in the hard task. Note also that results are on the response scale (with a value of 1 representing 100% accuracy).

**Estimated Marginal Means**

| | | | | 95% CI | |
|---|---|---|---|---|---|
| **P3_Centroid** | **Easy/Hard** | **Estimate** | **SE** | **Lower** | **Upper** |
| -28.333 | Easy | 0.959 | 0.006 | 0.945 | 0.969 |
| 16.861 | Easy | 0.968 | 0.005 | 0.957 | 0.976 |
| 62.054 | Easy | 0.975 | 0.005 | 0.964 | 0.983 |
| -28.333 | Hard | 0.761 | 0.024 | 0.711 | 0.805 |
| 16.861 | Hard | 0.765 | 0.024 | 0.716 | 0.808 |
| 62.054 | Hard | 0.769 | 0.026 | 0.714 | 0.816 |

**Replication Testing - Combined Analysis of the easy task data from both studies**

The current results differed from those reported in our previous study. The results of our previous study indicated there was a significant main effect of group in the TANOVA from -1 to 61ms, where meditators showed stronger occipital activation, and a significant main effect from group from 416 to 512ms, where meditators showed a more frontal P3 distribution. Our previous study results also showed an interaction between group and Go/Nogo in the GFP test from 336 to 449ms, where meditators showed more equal amplitudes of global neural response between Go and Nogo trials. To test whether the difference in results might be due to differences in EEG pre-processing, we performed an exploratory analysis of the data from our current study after applying the same subtraction baseline correction method as our previous paper (instead of the more appropriate regression baseline correction method implemented with our updated analysis pipeline). This analysis was also restricted to just the easy version of the task (in direct replication of our previous research). This analysis was intended to test whether the baseline correction method might have been the cause of the effects detected in our earlier study.

Interestingly, when implementing the subtraction baseline correction method the current study replicated the pre-C1 effect, with averaged data from -1 to 61ms showing a main effect of group ($p$ = 0.004, $np^2$ = 0.065, see Figure S4). The effect showed the same topographical difference between groups as our previous study, with meditators showing more positive posterior voltages and more negative fronto-central voltages than non-meditators. Since the subtraction baseline method transposes a mirror image of the topography from the baseline correction period into the active period, we suspect that instead of a true effect at -1 to 61ms post stimulus, the difference between groups in this period when the subtraction baseline correction method is applied is driven by a between group difference in the interaction between late positive potential / slow wave activity at the end of the previous trial (which is also the activity during the pre-stimulus baseline correction period) and activity in the -1 to 61ms period. Indeed, the between group difference in the distribution of activity is the mirror image of a main effect of group that was apparent in the regression baseline corrected analysis from 560 to 604ms, an effect that was significant in our primary analysis but did not pass the global duration controls) (see Figure S5).

However, unlike the effects within the pre-C1 period, no main effect of group was present in the TANOVA when activity was averaged from 416 to 512ms ($p$ = 0.770), and no interaction between group and Go/Nogo trial was present when activity was averaged from 336 to 449ms ($p$ = 0.392). This was the case even when data were high pass filtered at 0.1Hz in closer replication of our previous study (both $p > 0.05$).

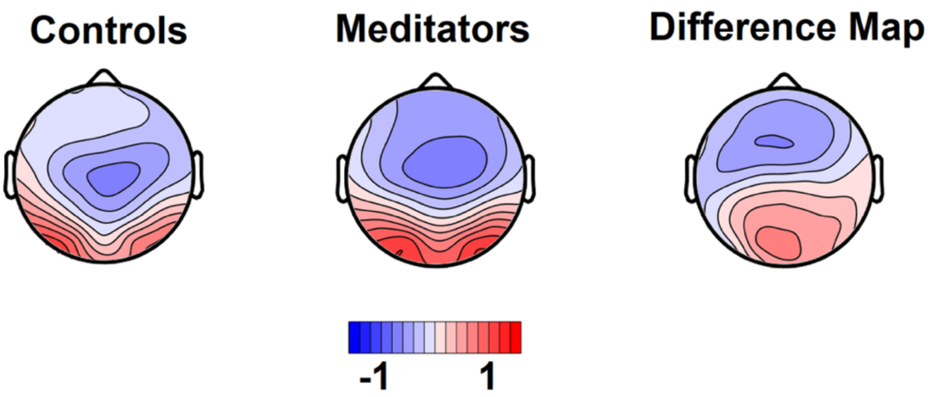


**Figure S4.** Topographical distributions during the pre-C1 period (-1 to 61ms) averaged across easy Go and Nogo trials, and a difference map reflecting meditator activity minus non-meditator activity.

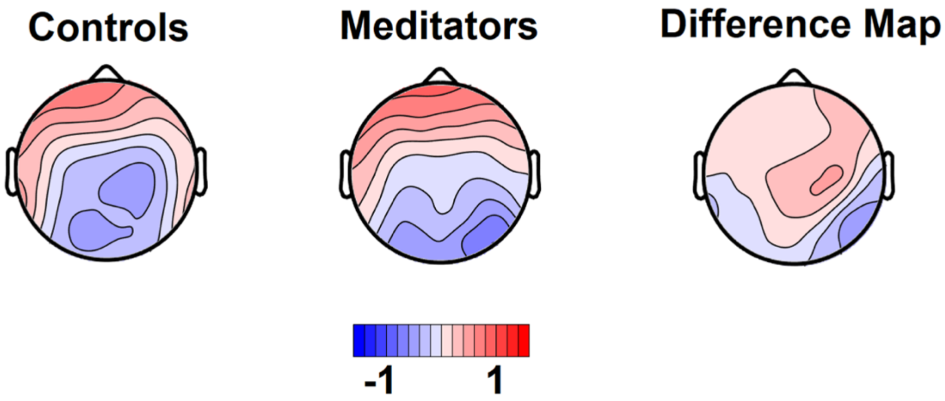


**Figure S5.** Topographical distributions during the baseline period (-100 to 0ms) averaged across easy Go and Nogo trials, and a difference map reflecting meditator activity minus non-meditator activity. Note that the maps are similar to an inverse of the pre-C1 period activity, indicating the effect a subtraction baseline comparison approach had on the pre-C1 effect detected in our previous study.

**Testing for differences in the topographical distribution of activity with the combined dataset**

To assess whether the increased power obtained from combining the two studies provided any additional effects, we re-processed the original study's data using our updated pre-processing pipeline and conducted an analysis using RAGU on the easy Go/Nogo task data. When the combined data from both studies were analysed, a main effect of group was present in the N2 window (from 191 to 244ms) which lasted longer than the global duration controls (global duration control = 49ms, $p$-value averaged across period of significance = 0.004, $np^2$ = 0.022, see Figure S6). This result aligns with the analysis of our age-matched sample in the current study, but we note the effect size is small.

Additionally, a main effect of group was present when data were averaged within the period of 416 to 512ms (where meditators showed significantly more frontal P3 activity in both trial types in our previous study). However, the effect was small, and only just exceeded the statistical threshold ($p$ = 0.047, $np^2$ = 0.002, see Figure S7). We also note that while the pattern was similar in our new study, the effect did not independently replicate with statistical significance in our new study data, it was only when the two studies were combined that the effect was significant. No other main effect or interaction between group and Go/Nogo trial type was significant (all $p > 0.05$).

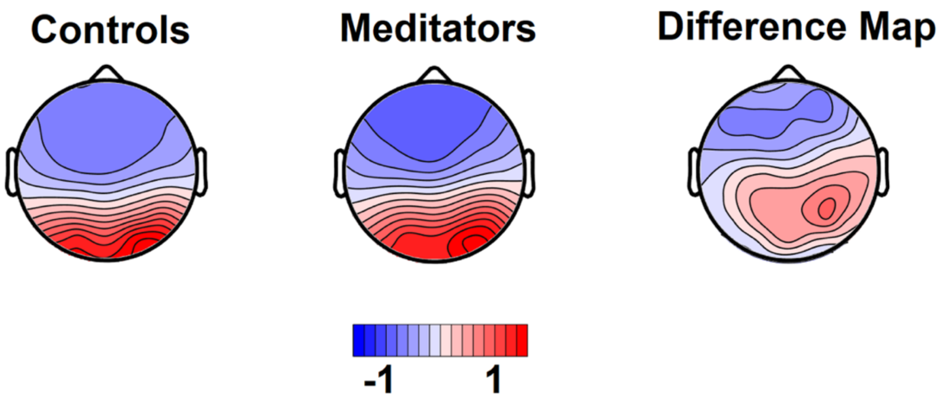


**Figure S6.** Mean N2 activity averaged within the significant period from the new study (198 to 241ms) and across both studies and the easy Go and Nogo trials, along with the difference map for meditator minus non-meditator (control) activity.

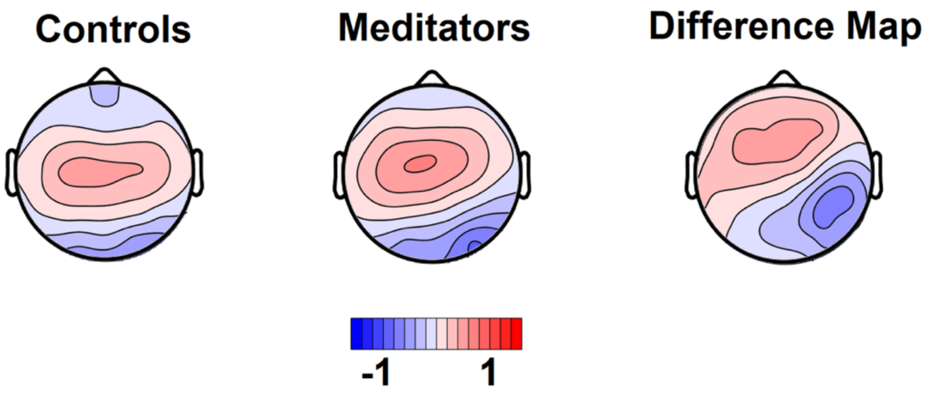


**Figure S7.** Mean N2 activity averaged within the significant group main effect TANOVA period from our old study (416 to 512ms) and across both studies and the easy Go and Nogo trials, along with the difference map for meditators minus control activity. Note that the effect size was very small, and only just exceeded the statistical threshold ($p$ = 0.047, $np^2$ = 0.002). We also note that the effect did not independently replicate in our new study data, it was only when the two studies were combined that the effect was apparent.

**Testing for differences in the global neural response strength with the combined dataset**

When the GFP was analysed using data from both studies, there was a significant interaction between group and Go/Nogo trial type in the P3 window (386 to 429ms) which lasted longer than global duration controls ($p$-value averaged across the period of significance = 0.007, $np^2$ = 0.062, global duration control = 43ms, see Figure S8). Post-hoc analysis of data averaged within this 386 to 429ms period indicated that meditators showed larger Nogo P3 GFP values than non-meditators ($p$ = 0.006, $np^2$ = 0.061), and that meditators also showed larger Nogo P3 GFP values than Go P3 GFP values ($p$ = 0.002, $np^2$ = 0.137), but there was no difference between groups for Go trials ($p$ = 0.752, $np^2$ = 0.001), and non-meditators did not show differences between their Go and Nogo trials ($p$ = 0.455, $np^2$ = 0.010).

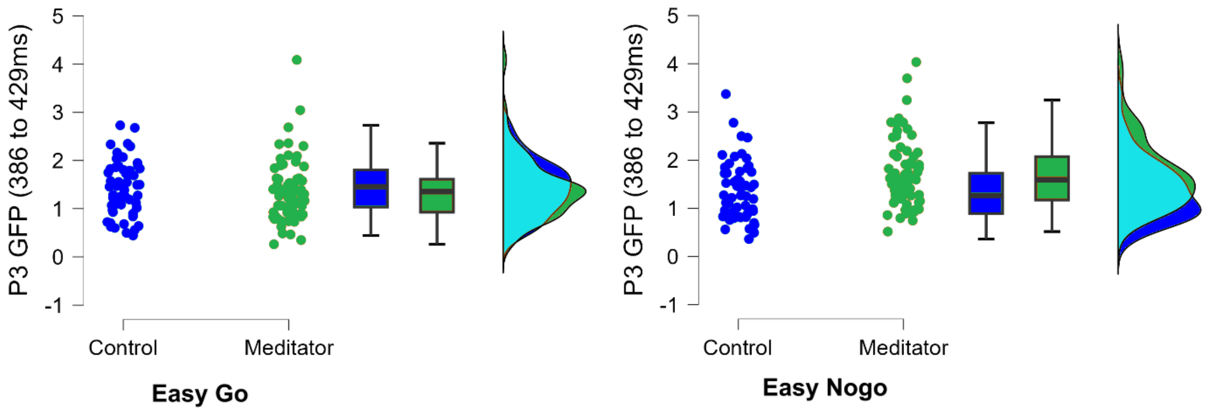


**Figure S8.** P3 amplitude averaged across the significant interaction period following responses to easy Go and Nogo stimuli across participants from both studies.

We note that this effect is within a similar window to the interaction in the GFP test between group and Go/Nogo trial type in our original study (336 to 449ms). However, in our previous study, the interaction suggested that non-meditators showed a larger Go P3 GFP than Nogo P3, while meditators did not show a difference between the conditions, which differs from the pattern shown in the current study. This result and pattern was consistent when we high pass filtered our data at 0.1Hz, indicating that the 0.5Hz high pass filter setting in the new study was not responsible for the altered effect compared to the original study ($p$-value averaged across the period of significance = 0.003, $np^2$ = 0.066). To test the reason for the discrepancy between our current results and our previous results, we also performed an analysis of this time window using a subtraction baseline correction approach. In this analysis, the result was non-significant ($p$ = 0.301, $np^2$ = 0.010). However, the analysis did show a similar pattern to the results reported in the original study, with meditators showing less difference between the Go and Nogo trials than controls. This suggests that our original result may have been driven by a combination of non-optimal data cleaning and the confounding influence of a subtraction baseline correction method, but was not produced by the filter settings.
